# DHX30 is a replication stress-responsive R-loop helicase that safe-guards genome stability

**DOI:** 10.64898/2026.10.25.761626

**Authors:** Elena R. Nedelcu, Nan Cher Yeo, Hannes Huber, Archana B. Prusty, Thinh Q. Truong, Nghi Dang, Sönke Harder, Johannes Lederbauer, Ivana Lessel, Nina Ripin, Christina Kiel, Utz Fischer, Hans-Jürgen Kreienkamp, Davor Lessel

## Abstract

R-loops are three-stranded nucleic acid structures formed primarily during transcription that can obstruct replication fork progression, leading to replication stress and genome instability. Although RNA helicases are increasingly implicated in R-loop metabolism, how individual helicases safeguard genome integrity remains incompletely understood. Here we identify DHX30, a neurodevelopmental disorder-associated DExH-box RNA helicase, as a replication stress-responsive regulator of R-loops. We show that DHX30 localizes to the nucleus in addition to its known cytoplasmic and mitochondrial pools, and that its nuclear abundance increases following aphidicolin or camptothecin treatment. DHX30 deficiency results in RNA:DNA hybrid accumulation in human cells and zebrafish, R-loop-dependent DNA double-strand breaks, micronucleus formation, reduced DNA synthesis, and altered expression of immediate-early stress genes under basal conditions and following replication perturbation. Biochemically, DHX30 binds branched nucleic acid structures and directly unwinds R-loop substrates containing a 3′ single-stranded RNA overhang in an ATP- and polarity-dependent manner. Proteomic analyses place DHX30 within a nuclear network of RNA- and R-loop-associated factors, including the RNA helicase DHX9. These findings establish DHX30 as a conserved R-loop-resolving helicase that protects from R-loop-induced replication perturbations, contributes to transcriptional homeostasis, and maintains genome stability, thereby providing a mechanistic framework for understanding *DHX30*-associated neurodevelopmental disorders.

## INTRODUCTION

RNA metabolism relies on the coordinated activity of RNA helicases, a large and diverse family of enzymes that participate in nearly all cellular processes involving RNA. Based on conserved sequence motifs and mechanistic features, RNA helicases are classified into six superfamilies (SF1–SF6) (1,2). Among these, the SF2 superfamily represents the largest and most extensively studied group. It includes helicases defined by conserved DExD or DExH motifs, giving rise to the DDX and DHX proteins, respectively (3,4). These helicases play central roles in RNA secondary structure remodelling and ribonucleoprotein (RNP) dynamics, and many transcriptional, post-transcriptional, and translational processes require the coordinated activity of multiple family members (5–7).

R-loops are three-stranded nucleic acid structures consisting of an RNA:DNA hybrid and a displaced single-stranded DNA. They are widespread across eukaryotic genomes and play regulatory roles in transcription, chromatin organization, and genome maintenance (8,9). However, excessive or aberrant R-loop accumulation can interfere with transcription and DNA replication, promote DNA damage, and drive genome instability (10,11). Notably, dysregulated R-loop metabolism has been implicated in neural development, where persistent R-loops can disrupt developmental gene programs and genome integrity, potentially contributing to neurodevelopmental disorders (12–14).

To maintain R-loop homeostasis, eukaryotic cells employ a multilayered surveillance system comprising DNA and RNA helicases, nucleases, RNases, chromatin-remodelling complexes, and ribonucleoproteins (RNPs) (15). Among these several DEAD- and DEAH-box helicases, including DDX1, DDX5, DDX17, and DHX9, have been shown to resolve R-loops and safeguard genome integrity (16–19). Despite these advances, how disease-associated RNA helicases that are not classically nuclear contribute to R-loop regulation remains poorly understood.

We previously identified rare germline variants in *DHX30*, encoding a DEAH-box RNA helicase, as the cause of a spectrum of neurodevelopmental disorders (20,21). DHX30 has been primarily characterized as a cytoplasmic and mitochondrial protein, however, its potential nuclear functions have remained unexplored. Here, we identify DHX30 as a replication stress-responsive nuclear R-loop helicase. DHX30 directly unwinds R-loop substrates containing a 3′ single-stranded RNA overhang, and its deficiency leads to RNA:DNA hybrid accumulation, DNA double-strand breaks, micronucleus formation, reduced DNA synthesis and altered stress-responsive gene expression, thereby linking defective R-loop resolution to *DHX30*-associated neurodevelopmental disorders.

## MATERIALS AND METHODS

### Cell culture

Human embryonic kidney 293T (HEK293T) and human bone osteosarcoma epithelial (U2OS) cells were grown on cell culture dishes and coverslips, respectively, utilizing Dulbecco’s Modified Eagle Medium (DMEM) supplemented with 10% fetal bovine serum (FBS). HEK293T medium additionally contained 5% pyruvate. Cells were maintained at 37°C in an incubator with 5% CO_2_.

### Cellular and Animal Models

Generation of a HEK293T DHX30 stable knockdown line and zebrafish DHX30 stable knockout has been described previously (20).

### Plasmid constructs

| **Plasmid** | **Reference/Supplier** |
| --- | --- |
| pet-28M-SUMO1::DHX30-WT | (20) |
| pet-28M-SUMO1::DHX30-R493H | (20) |
| pet-28M-SUMO1::DHX30-H562R | (20) |
| pEGFP-DHX30-cyto | (21) |
| pEGFP-DHX30-mito | (21) |
| pEGFP-RNASEH1 | Addgene (108699) |
| pCAG-DHX9-eGFP | (22) |

### Transfection and Treatments

The construct of the expression vectors for the DXH30 wild type have been described previously (21). HEK293T and U2OS cells were transfected for 24h with the transfection reagents TurboFect (ThermoFisher Scientific) or FuGene (Promega), respectively, according to the manufacturer’s recommendations. Transient DHX30 knockdown was performed using a synthetic siRNA targeting human DHX30 (QIAGEN; cat. no. SI04131911; reference transcript NM_014966; target sequence, 5′-CACGGGCAGCTGCTTGCGCTA-3′). AllStars Negative Control siRNA (QIAGEN; cat. no. 1027280), whose sequence is proprietary, served as the non-targeting control. Cells were transfected with siRNA at a final concentration of 50 nM using Lipofectamine RNAiMAX (Thermo Fisher Scientific), according to the manufacturer’s instructions, and analysed 24 h after transfection. The cells were treated with 2 µM camptothecin (CPT) (Sigma-Aldrich®) for 2h, with 5µg/ml actinomycin d (ActD) (Sigma-Aldrich®) for 8h, with 40 ng/ml Mitomycin C (MMC) (Sigma-Aldrich®) for 48h or with 2 µM APH (SigmaAldrich) for 24h.

### Antibodies

Primary antibodies used in this study: mouse anti-C23 (Nucleolin NCL; Santa Cruz Biotechnology, #sc-8031) used at a dilution of 1:50 for immunofluorescence experiments (IF); mouse anti-GFP Epitope Tag (BioLegend, #MMS-118P) used at a dilution of 1:5000 for western blotting experiments (WB); rabbit anti-GFP (abcam, #ab290) used at a dilution of 1:5000 for WB; rabbit polyclonal anti-DHX30 (Bethyl, #A302-218A) used at a dilution of 1:200 for IF and 1:2000 for WB; rabbit polyclonal anti-DHX9 (Bethyl, #A300-854A) used at a dilution of 1:200 for IF and 1:2000 for WB; mouse anti-gammaH2AX [p-Ser139] (Novusbio, #NB100-74435) used at a dilution of 1:200 for IF and 1:1000 for WB; rabbit anti-gammaH2AX [p-Ser139] (Millipore, #07-164) used at a dilution of 1:200 for IF; hFAB Rhodamine Anti-GAPDH Primary Antibody (Bio-Rad, #AbD22549) used at a dilution of 1:5000 for WB; chicken anti-GAPDH (Abcam, #ab83956) used at a dilution of 1:5000 for WB; goat anti-Lamin B1 (Sant Cruz Biotechnology, #sc-30264) used at a dilution of 1:2000 for WB; rabbit 53BP1 (Novus Biologicals, #NB100-305S) used at a dilution of 1:100 for IF; rabbit anti-RPA32/RPA2 phospho-S4/S8 (Abcam, #ab243866) used at a dilution of 1:100 for IF; and mouse anti-HSP90 (Proteintech, #60318-1) used at a dilution of 1:5000 for WB.

For detecting RNA:DNA hybrids, mouse anti-DNA-RNA Hybrid [S9.6] (Kerafast, #ENH001) 1:100 dilution for IF and 1:1000 dilution for dot blot.

### Immunofluorescence microscopy and EdU incorporation assay

Cells were grown on coverslips; after fixation with ice cold methanol, cells were washed with phosphate-buffered saline (PBS) and permeabilized for 5 min in 0.5% Triton X-100 in PBS. After blocking with 5% bovine serum albumin (BSA)/PBS for 1h, cells were incubated with primary antibody (as indicated in the antibody list), followed by washing in PBS and incubation with secondary antibody (Alexa Fluor 488/546/633, Invitrogen, anti-rabbit and anti-mouse respectively. To visualize the nucleus, cells were stained with 0.3 μg/ml DAPI. Slides were mounted using ProLong™ Diamond Antifade Mountant (Invitrogen) and images were acquired using confocal microscopes LSM700 (Zeiss) and Olympus FV3000 at 40X and 60X magnification. Images were analysed using Zen software (Zeiss) and Fiji. The Click-IT® Plus EdU imaging kit (Molecular Probes) was used to assess DNA synthesis assay following a 30-min EdU pulse immediately before fixation, according to the manufacturer’s recommendations, as previously described (23).

### Lysis, GFP-trap, Western blot and Cell fractionation analysis

Briefly, cells were harvested from an 80-90% confluent 10 cm plate and the pellet was resuspended in 0.5 mL lysis buffer (10 mM Tris-HCl, pH 7. 5, 150 mM NaCl, 2 mM EDTA, 1% TX-100, 10% glycerol, 1 mM DTT plus proteinase and phosphatase inhibitors) or immunoprecipitation buffer (IP buffer) (120 mM NaCl, 50 mM Tris-HCl, pH 8.0, 0.5% NP40, 1 mM EDTA plus proteinase and phosphatase inhibitors) and incubated for 20 min on ice. After centrifugation, the supernatant was added to GFP-Trap® magnetic agarose beads (ChromoTek) and GFP-Trap was performed according to the manufacturer’s recommendations. For nuclease treatment experiments, cell lysates were incubated with RNase A or RNase H for 30 min at 37 °C before GFP-Trap immunoprecipitation. For Coomassie staining of DHX30-associated proteins, RNase A digestion was performed in IP buffer containing 150 mM NaCl by adding 10 µL of RNase A stock solution (10 mg/mL) to 350 µL of lysate. For analysis of the DHX30-DHX9 association, lysates were mixed 1:1 with 3 M NaCl before RNase A treatment, yielding a final NaCl concentration of approximately 1.6 M. RNase H (BioLabs, #M0297L) treatment was performed under the manufacturer’s recommended conditions. Following nuclease treatment, GFP-Trap immunoprecipitation was performed as described above. For quantitative analysis of DHX30–DHX9 co-immunoprecipitation, the co-immunoprecipitated DHX9 signal was normalized to the corresponding recovered DHX30-GFP signal to account for differences in bait recovery between conditions. The GFP-Trap affinity purification used for mass spectrometry analysis was performed without nuclease treatment.

For Western blot, protein levels were measured using a BCA protocol. For protein separation, 8-12% polyacrylamide gels were used. After running, the gels were transferred to a nitrocellulose membrane. Incubation with antibodies and detection by chemiluminescence (ECL) were performed according to standard procedures.

Cellular fractionation was performed using the Nuclear/Cytosol Fractionation Kit (Abcam, #ab289882). Fractionation was assessed by immunoblotting using Lamin B1 as a nuclear marker and GAPDH and HSP90 as cytoplasmic markers.

### Dot Blot and Slot Blot analyses

Following manufacturer’s instructions, total DNA was purified using QIAamp DNA Blood Kits - Genomic DNA Extraction DNA (Qiagen, #51104). Concentration was determined using a spectrophotometer. To determine the R-loop content, the DNA was blotted on a positive charged nylon transfer membrane (Amersham), prewetted with 10X SSC, using a Bio-Dot/Slot Apparatus (Bio-Rad) in duplicate. The membrane was UV crosslinked (120 mJ/cm2) and probed with S9.6 antibody (Kerafast, #ENH001; 1:5000) in TBS-T solution. To test the specificity of S9.6 antibody, samples were additionally digested with RNaseH (NEB). After incubations in S9.6 antibody, blots were incubated with HRP-conjugated secondary antibodies and S9.6 signals were visualized using a ChemiDoc (Bio-Rad). Images were analysed using Quantity One software (Bio-Rad). Methylene blue (Sigma-Aldrich, #M4159) staining was used to assess total DNA loading, and S9.6 signal intensities were quantified and normalized to the corresponding methylene blue signal.

### Neutral Comet Assay

Neutral comet assay was performed using the CometAssay® Kit (R&D Systems, Cat. No. 4250-050-K) according to the manufacturer’s instructions, using 1xTBE buffer for neutral electrophoresis. Cells were processed and embedded in LMAgarose as specified by the manufacturer, fol-lowed by lysis and neutral electrophoresis in 1× TBE for 40 min at a field strength of 1 V/cm.DNA was stained with DAPI. Fluorescence images were acquired using a KEYENCE BZ-X800 fluorescence microscope equipped with a 20x objective, using identical acquisition settings for each biological repeat condition. Images were analysed in Fiji/ImageJ using OpenComet v1.3.1. Comet identification was manually reviewed, and only individual, non-overlapping, correctly segmented comets were included in the analysis. Four independent biological experiments were performed, with a total of 500 comets pooled across the four experiments analysed. DNA break-associated comet parameters were quantified as Tail DNA (%), Olive tail moment, and tail length.

### Micronucleus assay

Proliferating cells were stained with DAPI, and images were acquired using a KEYENCE BZ-X800 fluorescence microscope equipped with a 20x objective. The total number of nuclei and the number of nuclei with associated micronuclei were counted using ImageJ, as previously described (24). Micronucleus frequency was calculated as the percentage of nuclei with associated micronuclei. At least 300 cells were scored per biological replicate, with more than 1,000 cells analysed in total across three independent biological replicates.

### RNA-sequencing analysis

For bulk RNA-sequencing, total RNA was extracted using the EZ2 Connect MDx instrument (Qiagen) with the kit EZ2 RNA/miRNA Tissue/Cells Kit (Qiagen), and RNA concentration and quality was estimated using a spectrophotometer. mRNA selection through poly(A) capture and library preparation was performed using the Stranded mRNA Prep, Ligation kit (Illumina). All libraries were sequenced by a NextSeq 550 System (Illumina) generating 149 bp paired end reads. Quality control was performed during all steps of analysis by applying FastQC v.0.12.1 (https://www.bioinformatics.babraham.ac.uk/projects/fastqc/) and MultiQC v.1.25.2. (25). Trimming of raw reads was performed using fastp v.0.24.0 (26), including detection of paired-end adapters, read length requirement of 36 bp, trimming of one bp at the tail of the read and low complexity filter with default settings. Reads were aligned to the human reference genome GRCh38 primary assembly and gene annotation v48 obtained from GENCODE (27) using the STAR aligner v.2.7.10b (28) in two-pass mode and by using ENCODE standard option, except the maximum number of mismatches per pair relative to the read length argument which was kept as default. Gene-level expression counts were estimated using RSEM toolbox v.1.2.28 (29) with standard parameters. For the principal component analysis (PCA), counts were calculated as Counts per Million (CPM) by applying the *cpm* function implemented in the edgeR package v.4.4.0 (30,31) in R v.4.4.2. (https://www.R-project.org/). CPM values were log2 transformed and the PCA was performed with the *prcomp* function implemented in R. To detect differentially expressed genes the classical edgeR pipeline was used (31). Briefly, genes with low expression were filtered out using the *filterByExpr* function. Normalization was performed using the trimmed mean of M-values (TMM) method with the *normLibSizes* function. The estimate common and tagwise dispersion was calculated using the *estimateDisp* function and the *exactTest* function was applied to obtain differentially expressed genes. Genes were differentially expressed with a false discovery rate (FDR) below 0.001 and an absolute log fold change (FC) greater than 1. For pathway enrichment analysis ENSEMBL IDs obtained from GENCODE were converted to Entrez IDs using the biomaRt package v.2.62.0 (30). GO Biological Process enrichment analysis was conducted using the *enrichGO* function from clusterProfiler (v.4.14.0) (32). Pathways were considered significantly enriched at an adjusted p-value below 0.05. Volcano plots and boxplots were generated using the ggplot2 package v.3.5.1 (Wickham, H. (2016) Elegant graphics for data analysis). Heatmaps were created using the pheatmap package v.1.0.12 (https://CRAN.R-project.org/package=pheatmap).

### Reverse transcription and quantitative PCR

Total RNA (1 µg) was combined with 1 µL of random hexamer primers in a volume of 12 µL and incubated at 65 °C for 5 min. Subsequently, 1 µL of reverse transcriptase, 4 µL of 5x reaction buffer, 2 µL of dNTPs and 1 µL of nuclease-free water were added to a final volume of 20 µL. Reverse transcription was performed at 25 °C for 10 min, 42 °C for 60 min and 70 °C for 10 min. Quantitative PCR was performed using TaqMan-based detection on a QuantStudio Real-Time PCR System (Applied Biosystems). Each 10-µL reaction contained 5 µL of 2× TaqMan Master Mix, 1 µL each of forward and reverse primers, 0.125 µL of the corresponding fluorescent probe, 0.375 µL of nuclease-free water and 2.5 µL of diluted cDNA. Reactions were performed in technical triplicates. Cycling conditions comprised 95 °C for 10 min, followed by 40 cycles of 95 °C for 15 s and 60 °C for 1 min, with fluorescence acquisition during the latter step. Relative expression was normalized to HPRT1 and calculated using the 2 method. Primer and probe sequences are provided in Supplementary Table 1.

### Mass Spectrometry

GFP-tagged DHX30 or GFP alone was affinity-purified from HEK293T nuclear extracts using GFP-Trap_A matrix (Chromotek), without nuclease pretreatment, as previously described (20). Briefly, proteins were denatured in 100 µL Laemmli buffer. Disulfide bonds were reduced in 10 mM dithiothreitol for 30 min at 56 °C and alkylated in presence of 20 mM iodoacetamide for 30 min at 37 °C in the dark. Acetonitrile (ACN) was added to a concentration of 70% and 1 µL carboxylate modified magnetic beads (GE Healthcare Sera-Mag™, Chicago, USA) at 1:1 (hydrophilic/hydrophobic) ratio in methanol/LC-MS grade water were added following the single-pot, solid-phase enhanced sample preparation (SP3)-protocol workflow. Samples were shaken at 1400 rpm for 18 min at room temperature. Tubes were placed on a magnetic rack, and the supernatant was removed. Magnetic beads were washed two times with 100% ACN and two times with 70% Ethanol on the magnetic rack. After resuspension in 50 mM NaHCO_3_, digestion with 1 µg trypsin (sequencing grade, Promega) was performed at 37°C overnight while shaking at 1400 rpm. Tubes were placed on the magnetic rack, the supernatant was removed, and the beads were washed twice with 100% ACN. Elution was performed with 2% DMSO in 1% formic acid (FA). The supernatant was dried in a vacuum centrifuge and stored at −20°C until further use. Chromatographic separation of peptides was achieved with a two-buffer system (buffer A: 0.1% FA in H2O, buffer B: 0.1% FA in ACN) on a nano-UHPLC (Dionex Ultimate 3000 UHPLC system, Thermo Fisher). Attached to the UHPLC was a peptide trap (100 µm x 20 mm, 100 Å pore size, 5 µm particle size, C18, Nano Viper, Thermo Fisher) for online desalting and purification, followed by a 25 cm C18 reversed-phase column (75 µm x 250 mm, 130 Å pore size, 1.7 µm particle size, peptide BEH C18, nanoEase, Waters). Peptides were separated using an 80 min method with linearly increasing ACN concentration from 2% to 30% ACN over 60 minutes. MS/MS measurements were performed on a quadrupole-orbitrap hybrid mass spectrometer (QExactive, Thermo Fisher Scientific). Eluting peptides were ionized using a nano-electrospray ionization source (nano-ESI) with a spray voltage of 1,800 V and analyzed in data dependent acquisition (DDA) mode. For each MS1 scan, ions were accumulated for a maximum of 240 milliseconds or until a charge density of 1 x 106 ions (Automatic Gain control or AGC target) was reached. Fourier-transformation based mass analysis of the data from the orbitrap mass analyzer was performed covering a mass range of m/z 400 – 1,200 with a resolution of 70,000 at m/z = 200. Peptides being responsible for the 15 highest signal intensities per precursor scan with an intensity threshold of 5 x 103 and charge state from +2 to +5 were isolated within a m/z 2 isolation window and fragmented with a normalized collision energy of 25% using higher energy collisional dissociation (HCD). MS2 scanning was performed, covering a mass range starting at m/z 100 and accumulated for 50 ms or to an AGC target of 1 x 105 at a resolution of 17,500 at m/z = 200. Already fragmented peptides were excluded for 20 s. With Proteome Discoverer 3.1.0.638 (Thermo Scientific, Bremen, Germany) the LC-MS/MS data were processed. Identification of the proteins from the MS/MS spectra were performed with the search engine Chimerys HT using the *Homo sapiens* SwissProt database (www.uniprot.org) and a contaminant database. For the searches the following parameters were applied: Precursor mass tolerance: 10 ppm. Fragment mass tolerance: 0.2 Da. Two missed cleavages were allowed. Carbamidomethylation on cysteine residues as a fixed modification and oxidation of methionine residues as a variable modification was used for the search. Peptides with a FDR of 1% using Percolator were identified. At least two unique peptides per protein were used as a condition for a reliable identification.

### Protein purification

pET-28M-SUMO1:DHX30 vector wild-type and mutant were obtained previously (20). Briefly, full-length DHX30 wild-type and mutant proteins were expressed in E. coli BL21 (DE3) Rosetta competent cells grown at 37 °C in LB broth supplemented with kanamycin until reaching an OD600 of 0.6–0.7. Protein expression was then induced by the addition of 0.5 mM IPTG, and the cultures were incubated overnight at 16 °C with shaking at 180 rpm. Cells were harvested by centrifugation and resuspended in lysis buffer (40 mM HEPES, 800 mM NaCl, 10 mM imidazole, 0.5 mM DTT, AEBSF and cOomplete protease inhibitors). Cells were lysed on ice, using three rounds of sonication for 2 min (1 s pulse, 1 s pause) at 60% amplitude, output control of 7, followed by 1 min break using a Branson 450 sonifier. The samples were clarified by centrifugation for 30 min at 20000 rpm and 4 °C, and the cleared supernatant was filtered through a 0.45 µm syringe filter and loaded onto pre-equilibrated prepacked HisTrap FF column (Cytiva). After washing, the protein was eluted with elution buffer (40 mM HEPES; 800 mM NaCl; 255 mM imidazole; 0.5 mM DTT; protease inhibitors). The eluted protein was dialyzed against 3 L dialysis buffer (20 mM HEPES; 800 mM NaCl, 1 mM DTT, 1 mM EDTA, 5 % Glycerol, protease inhibitors) over night, filtered through a 0.2 µm syringe filter and subsequently concentrated using an Amicon® Ultra Zentrifugenfilter, 30 kDa MWCOand subjected to further purification by size-exclusion chromatography using a Superdex 200 High Load column on an Äkta™ Purifier system (GE Healthcare) in 20 mM HEPES; 800 mM NaCl, 1 mM DTT, 5 % Glycerol and protease inhibitors.

### Preparation of R-loop and D-loop substrates

50 pmol of DNA or RNA oligonucleotides (Supplementary Table 2) were 5′-end radiolabelled with γ-[^32P]ATP using T4 polynucleotide kinase (Thermo Fisher Scientific) according to the manufacturer’s instructions in a 20 μl reaction. Depending on the desired overhang, oligonucleotides R3, R4, or R5 were used. After incubation for 1 h at 37°C, unincorporated nucleotides were removed using a G-25 spin column (1,000 × g, 5 min). The radiolabelled oligonucleotide was mixed with complementary DNA oligonucleotide B11 at a 1:2 molar ratio in 1× SSC buffer supplemented with 10 mM MgCl₂. Samples were heated to 95°C for 5 min and allowed to cool gradually to room temperature to permit annealing. Annealed RNA–DNA or DNA–DNA duplexes were resolved on an 8% native polyacrylamide gel in 0.5× TBE buffer. Electrophoresis was performed until the bromophenol blue dye migrated to approximately half the length of the gel. The desired duplex bands were excised and extracted overnight at room temperature in RNA extraction buffer (200 mM Tris-HCl, pH 7.0, 0.1% SDS, 1 mM EDTA) with gentle agitation. Duplexes were precipitated with one volume of isopropanol and 1 μl glycogen at −20°C for 1 h, centrifuged at 12,000 × g for 45 min, washed with 70% ethanol, air-dried, and resuspended in 1× SSC buffer containing 10 mM MgCl₂. To generate partial R-loop substrates, 50 pmol of partially complementary DNA strand (oligonucleotide T1) were added to the annealed duplexes. Samples were heated gradually to 65°C, incubated for 15 min, and then allowed to cool slowly to room temperature. The final annealed structures were resolved on an 8% native polyacrylamide gel in 0.5× TBE, excised, extracted as described above, and stored in 1× SSC buffer supplemented with 10 mM MgCl₂.

FAM-labelled and unlabelled RNA and DNA oligonucleotides (Supplementary Table 3) were purchased from commercial suppliers and used without further purification. Depending on the desired nucleic acid structure, oligonucleotides were mixed at a molar ratio of 1:1.5 (labelled: unlabelled) in annealing Buffer (AB) consisting of 1× SSC (sodium chloride and sodium citrate) supplemented with 10 mM MgCl₂. Samples were heated to 95 °C for 5 min in a heating block, which was then switched off to allow gradual cooling to room temperature and strand annealing.

### Helicase activity assay

Unwinding reactions were performed in a 20 μl reaction mixture containing up to 40 nM purified protein, 25 fmol radiolabelled nucleic acid substrate, 17 mM HEPES-KOH (pH 7.5), 150 mM NaCl, 1 mM MgCl₂, 2 mM DTT, 1 mM spermidine, 0.3% PEG 8000, 5% glycerol, 20 U RNasin™ Plus RNase inhibitor (Promega), and 1 mM ATP. Reactions were incubated for 30 min at 37°C and stopped by addition of 2X non-denaturing loading dye. Products were resolved by native polyacrylamide gel electrophoresis (8% gel; 19:1 acrylamide: bisacrylamide) in 0.5X TBE at 4°C. Reaction products were visualized by autoradiography.

### Electrophoretic Mobility Shift Assay (EMSA)

For electrophoretic mobility shift assays (EMSA), DHX30 (80 nM) was incubated with 2 nM of the indicated fluorescent nucleic acid substrates (listed in Supplementary Table 3) for 30 min at 37 °C. Reactions were performed in a buffer containing 100 mM HEPES, 250 mM NaCl, 250 mM KCl, 10 mM MgCl₂, 5 mM DTT, and 5% (v/v) glycerol. Samples were loaded onto 8% native polyacrylamide gels and resolved by electrophoresis in 1X TBE buffer. Gels were subsequently imaged for fluorescence using a ChemiDoc imaging system equipped with a blue-light transilluminator.

### Statistical analysis

Statistical analyses were performed using GraphPad Prism version 10.4.2. Data are presented as mean ± SEM unless otherwise indicated. All experiments were performed using independent biological replicates. The numbers of biological replicates, statistical tests and multiple-comparison procedures are specified in the corresponding figure legends. Where individual cells, nuclei or comets are displayed, the numbers analysed are also indicated. All t-tests were two-tailed. A *P* value < 0.05 was considered statistically significant. For RNA-seq and enrichment analyses, significance thresholds were based on adjusted *P* values or false discovery rates, as described in the respective Methods sections.

## RESULTS

### DHX30 localizes to the nucleus and its nuclear abundance increases upon replication stress

Previous studies have primarily described DHX30 as a cytoplasmic and mitochondrial protein in mammalian cells (21,33,34). However, careful re-examination of endogenous DHX30 by immunofluorescence microscopy in HEK293T and U2OS cells revealed, in addition to the expected cytoplasmic signal, a distinct nuclear signal overlapping with DAPI-stained regions (Fig. 1A and Supplementary Movie 1). Consistent with this observation, overexpression of a cytoplasmic GFP-tagged DHX30 isoform also resulted in detectable nuclear localization. In comparison, however, overexpression of a mitochondrial GFP-tagged DHX30 isoform resulted in only faint nuclear localization (Supplementary Fig. S1). To validate these findings using an orthogonal approach, we performed cytoplasmic and nuclear fractionation followed by immunoblot analysis. Fractionation efficiency was independently validated using Lamin B1 as a nuclear marker and HSP90, together with GAPDH, as cytoplasmic markers (Supplementary Fig. S2b). Again, DHX30 was detected in the cytoplasmic fraction, but was also clearly detectable in the nuclear fraction (Fig. 1B and C).

**Fig. 1.**
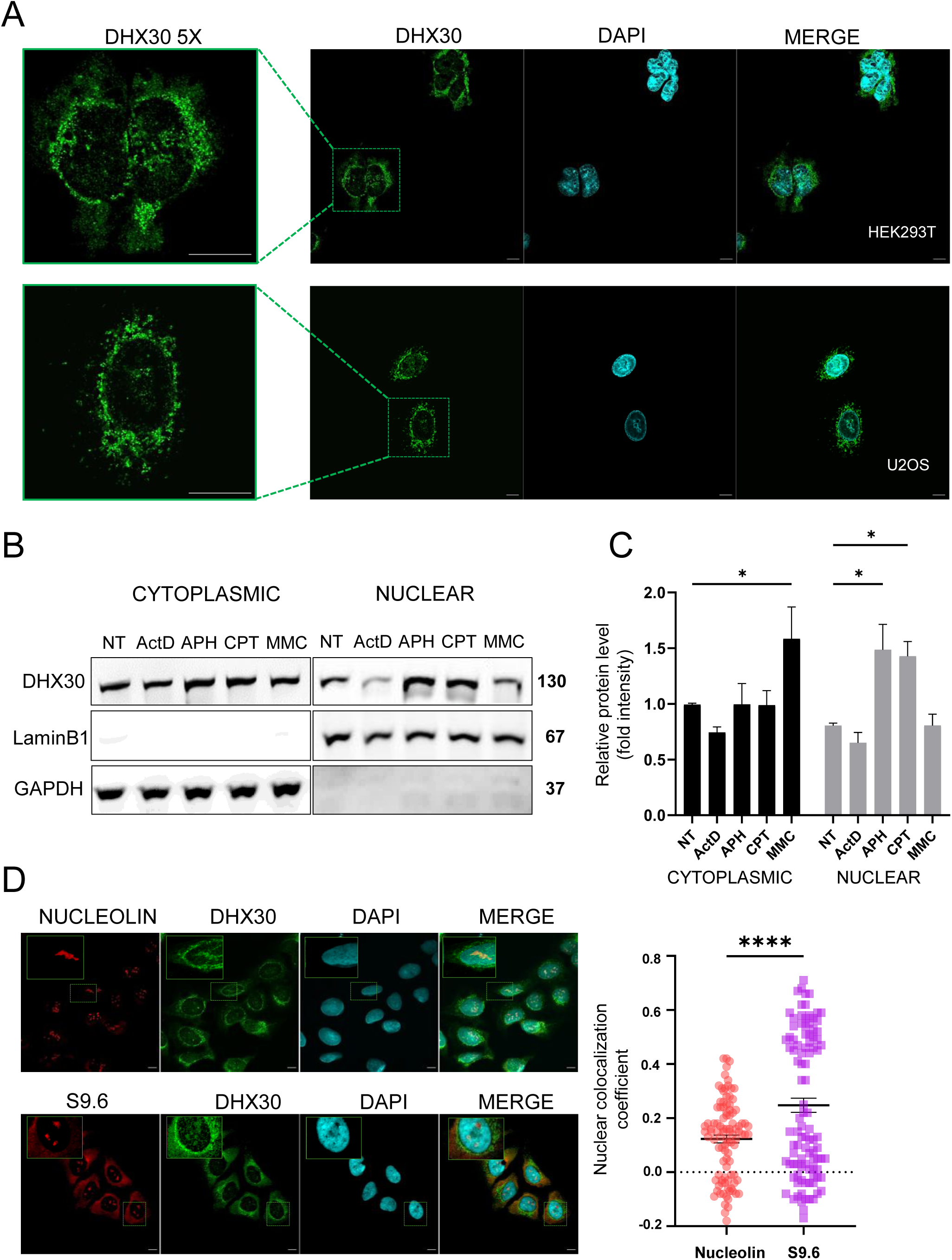
Nuclear DHX30 partially colocalizes with S9.6-positive regions and its nuclear fraction increases following replication stress. (A) Confocal imaging of HEK293T and U2OS cells stained for endogenous DHX30 (green) and nuclear marker DAPI (cyan). Higher-magnification views of the indicated regions are shown on the left, highlighting DHX30 localisation in both the nucleus and cytoplasm. (B) Immunoblot analysis of cytoplasmic and nuclear fractions isolated from HEK293T cells under untreated (NT) conditions or following exposure to ActD 5 µg/mL/8 h; APH 2 µM/24 h; CPT 2 µM/2 h; MMC 40 ng/mL/48 h. GAPDH and Lamin B1 were used as cytoplasmic and nuclear markers, respectively. (C) Quantification of DHX30 levels in cytoplasmic and nuclear fractions from (B). Data are presented as mean ± SEM from three independent biological experiments. Statistical significance was assessed using two-way ANOVA followed by Dunnett’s multiple comparison test comparing each treatment with the corresponding untreated (NT) condition within each cellular fraction (*P < 0.05). (D) Representative single-plane immunofluorescence images of U2OS cells co-stained with DHX30 ab (green) and either nucleolin ab (red) or S9.6 ab (red) and DAPI (cyan). Insets highlight regions of nuclear enrichment. Scale bar, 10 μm. On the right colocalization study of DHX30 with S9.6 or nucleolin. Nuclear colocalization coefficient was quantified using Pearson’s R (no threshold) calculated with Coloc2 (Fiji). Each point represents one cell. A total of 100 cells pooled from three independent biological experiments were analyzed. Statistical analysis was performed using unpaired T-test, two-tailed (**** P < 0.0001).

Given that some RNA-binding proteins (RBPs) undergo stress-dependent changes in sub-cellular localization (35,36), we next investigated whether DHX30 subcellular localization is regulated in a similar manner. HEK293T cells were exposed to a panel of well-characterized stressors inducing DNA damage or transcriptional perturbations via distinct mechanisms: aphidicolin (APH), a DNA polymerase inhibitor; camptothecin (CPT), a topoisomerase I inhibitor; mitomycin C (MMC), a DNA cross-linking agent; and actinomycin D (ActD), a transcriptional inhibitor. Notably, DHX30 nuclear abundance increased specifically following APH and CPT treatment, two agents that induce replication stress and are associated with elevated transcription-replication conflicts and R-loop accumulation (37,38). In contrast, neither transcriptional inhibition by ActD nor DNA cross-linking by MMC resulted in detectable changes in DHX30 nuclear abundance. These findings demonstrate that DHX30 undergoes selective nuclear enrichment in response to replication stress, distinguishing it from a general DNA damage or transcriptional stress response (Fig. 1C).

Immunofluorescence analysis revealed that DHX30 is not uniformly distributed throughout the nucleus, but instead exhibits a distinct subnuclear, compartment-like localization pattern. To further characterize this distribution in more detail, we performed colocalization analyses using antibodies against DHX30 in combination with either nucleolin, a marker of the nucleolus, or S9.6, which recognizes RNA:DNA hybrids but can also bind double-stranded RNA (dsRNA) (39). While DHX30 showed partial overlap with nucleolin, it displayed greater spatial overlap with S9.6-positive nuclear signal (Fig. 1D).

Given that R-loops accumulate under conditions of replication stress and that DHX30 nuclear abundance increases following APH and CPT treatment, we next examined whether DHX30-S9.6-association is enhanced under these conditions. Confocal microscopy confirmed a robust increase in S9.6 signal intensity following treatment with either APH or CPT (Supplementary Fig. S3a and b). Notably, quantitative colocalization analysis revealed a significant increase in the overlap coefficient between DHX30 and S9.6 signals in cells exposed to these replication stressors (Supplementary Fig. S3c). Together, these data and the previous identification of DHX30 in RNA:DNA hybrid-associated proteomes (40,41) prompted us to investigate its potential role in R-loop regulation.

### DHX30 deficiency increases RNA:DNA hybrid abundance in human cells and zebrafish brains

We next investigated whether DHX30 contributes to RNA:DNA hybrid homeostasis. To this end, we analysed RNA:DNA hybrid abundance in previously established DHX30-deficient HEK293T cells, with a residual DHX30 immunoreactivity (20). Using S9.6-based dot blot analysis, we observed a significant increase in RNA:DNA hybrid signal in DHX30-deficient cells compared with wild-type HEK293T controls (Fig. 2A and B). RNase H treatment markedly reduced the signal, supporting its specificity for RNA:DNA hybrids (Fig. 2A). The increase was also observed after normalization to methylene blue staining of total DNA in additional dot blot experiments (Supplementary Fig. S4). Consistent with these findings, siRNA-mediated depletion of DHX30 in U2OS cells, with a residual DHX30 immunoreactivity of ∼30% (Supplementary Fig. S5a), led to increased S9.6-positive nuclear signal (Supplementary Fig. S5b and c). Quantitative analysis revealed significantly increased nuclear S9.6 intensity under basal conditions, which persisted and, in some cases, became more pronounced upon replication stress induced by APH or CPT (Supplementary Fig. S5b and c). These results indicate increased susceptibility to replication stress in the absence of normal DHX30 levels.

**Fig. 2.**
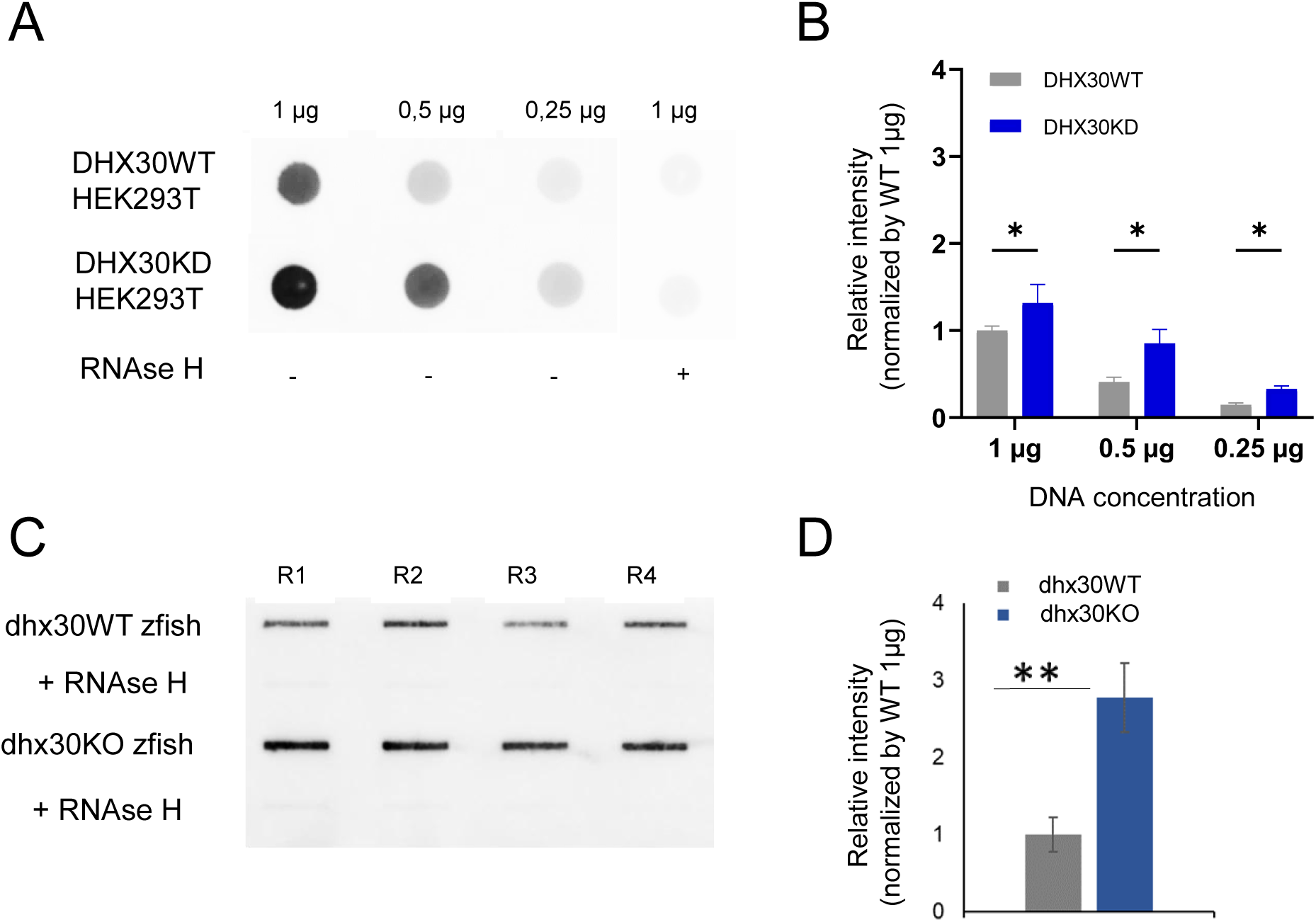
DHX30 deficiency leads to increased RNA:DNA hybrid accumulation in human cells and zebrafish brain. (**A**) Dot blot analysis of RNA:DNA hybrids detected with the S9.6 antibody in genomic DNA isolated from HEK293T-WT and DHX30KD cells. Equal amounts of DNA (1 μg, 0.5 μg and 0.25 μg) were spotted onto the membrane. *In vitro* RNase H treatment was used as a specificity control for RNA:DNA hybrids. (**B**) Quantification of S9.6 signal intensity normalized to WT 1µg signal. Data are presented as mean ± SEM; n =3. Statistical analysis was performed using two-way ANOVA followed by Holm-Sidak’s multiple comparison test (* P < 0.05). (**C**) RNA:DNA hybrids slot blot analysis of genomic DNA isolated from the brains of DHX30 knockout zebrafish and wild-type. Membranes were probed as described above. (**D**) Quantification of four biological replicates using BioRad Image Lab software. Mean of each experimental group was statistically analysed using t-test (**P < 0.005).

To determine whether this function of DHX30 is evolutionarily conserved, we next examined RNA:DNA hybrid abundance in a previously described *dhx30* homozygous mutant zebrafish model (20). Slot blot analysis of nucleic acid extracts prepared from zebrafish brain tissues similarly revealed elevated RNA:DNA hybrid signal in *dhx30* mutant animals relative to wild-type controls (Fig. 2C and D). Importantly, in both HEK293T cells and zebrafish samples, RNase H treatment markedly reduced this signal, supporting that the detected signals correspond to bona fide RNA:DNA hybrids (Fig, 2A and C).

Collectively, these findings show that DHX30 plays an active role in limiting RNA:DNA hybrid accumulation in both human cells and zebrafish, establishing an evolutionarily conserved function for DHX30.

### DHX30 helicase preferentially unwinds R-loop substrates containing a 3′ single-stranded RNA overhang

We have previously shown that DHX30 is an ATP-dependent helicase capable of unwinding RNA:RNA duplexes (20). Given its spatial overlap with S9.6-positive nuclear regions and the accumulation of RNA:DNA hybrids upon DHX30 depletion, we next asked whether DHX30 can directly process defined R-loop substrates. To address this, His₆–SUMO-tagged wild-type DHX30 was expressed in E. coli Rosetta cells and purified to near homogeneity (Supplementary Fig. S6). We first examined FAM-labelled substrate binding using electrophoretic mobility shift assays (EMSA). Purified DHX30 efficiently bound both R-loop and D-loop substrates, whereas no detectable binding was observed to double-stranded RNA:DNA hybrid (Supplementary Fig. S7). These data suggest preferential binding to the tested three-stranded substrates over the simple RNA:DNA duplex under these assay conditions.

We next assessed the helicase activity of DHX30 using radiolabelled substrates assembled in defined configurations: R-loop and D-loop substrates bearing a 3’ overhang, R-loop and D-loop substrates bearing a 5’ overhang, blunt-ended R-loop and D-loop substrates. Proper formation of R-loop substrates was verified by RNase H digestion *in vitro* (Supplementary Fig. S8). Increasing concentrations of recombinant protein were incubated with equimolar amounts of each substrate and the resulting unwinding products were resolved by native polyacrylamide gel electrophoresis (Supplementary Fig. S8b-d). DHX30 preferentially unwound R-loop substrates containing a 3′ single-stranded RNA overhang, consistent with the requirement of many DExH-box helicases for a 3′ single-stranded region to enable efficient loading and strand separation (Fig. 3A-C) (7,42,43).

**Fig. 3.**
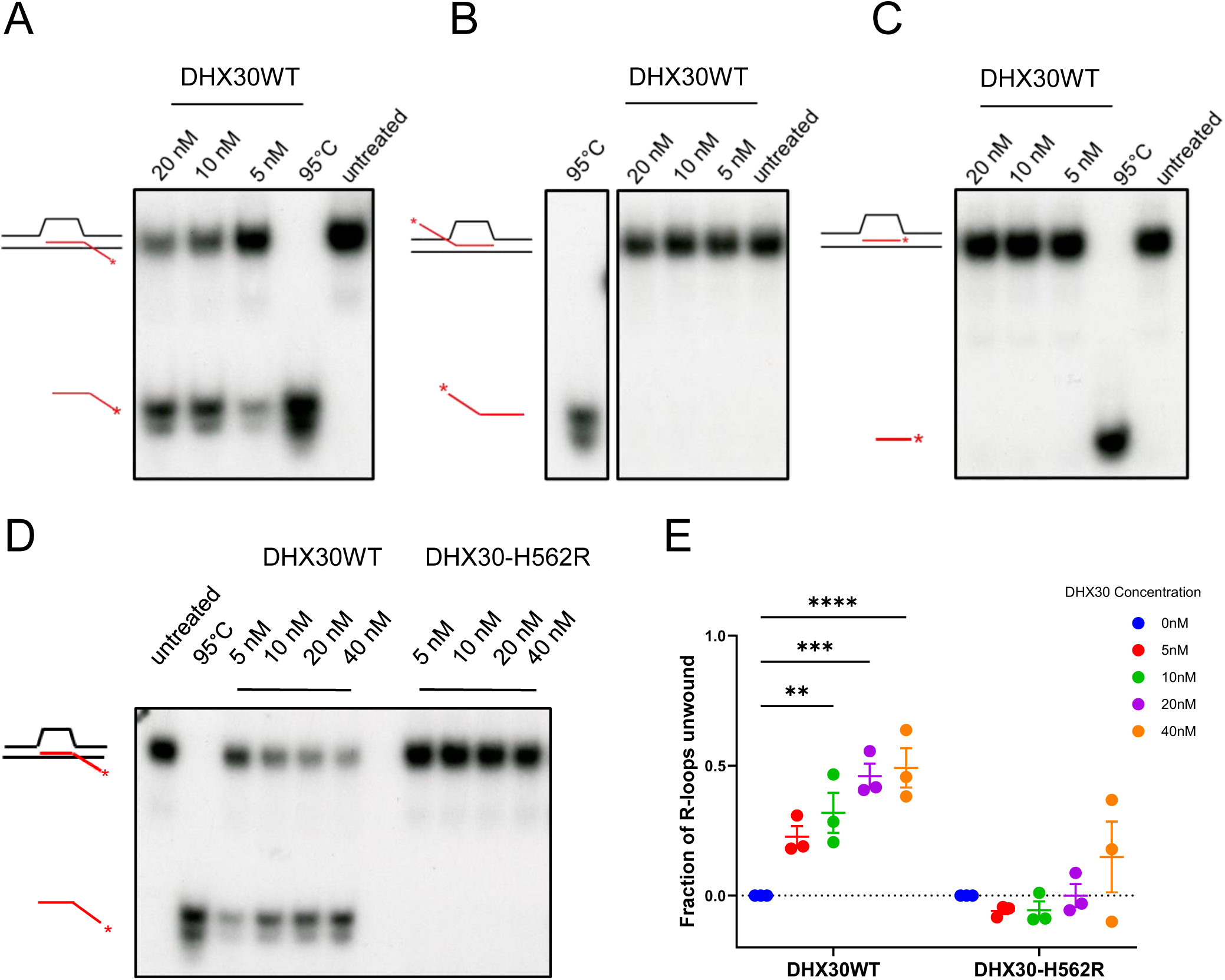
DHX30 helicase preferentially processes R-loop substrates containing a 3′ overhang. (A-C) Representative native PAGE analysis of R-loop unwinding by recombinant wild-type DHX30 (DHX30WT) using defined R-loop substrates containing a 3 ′ single-stranded RNA overhang **(A),** a 5′ single-stranded RNA overhang **(B),** or blunt ends **(C).** Reactions were performed with the indicated concentrations of DHX30WT. Heat-denatured (95 °C) and untreated substrates were included as controls. Schematics indicate the corresponding R-loop substrates; the red line represents the radiolabelled RNA strand and the asterisk denotes the radiolabelled end. Positions of intact R-loop substrate and released radiolabelled ssRNA are indicated. **(D)** Representative native PAGE analysis comparing unwinding of the 3′ single-stranded RNA overhang R-loop substrate by DHX30WT and the disease-associated DHX30-H562R variant at the indicated protein concentrations. **(E)** Quantification of R-loop unwinding by DHX30WT and DHX30-H562R shown as the fraction of substrate unwound at the indicated protein concentrations. Data are presented as mean ± SEM from three independent experiments. Statistical significance was assessed using two-way ANOVA followed by Dunnett’s multiple-comparisons test, comparing each protein concentration with the corresponding untreated condition within DHX30WT and DHX30-H562R (**P < 0.01; ***P < 0.001; **** P < 0.0001).

To further assess the specificity of DHX30-mediated R-loop unwinding, the 3′ overhang substrate was incubated with two disease-associated DHX30 variants, H562R and R493H, which were previously shown to be defective in RNA duplex unwinding (20). In contrast to wild-type DHX30, H562R showed nodetectable R-loop unwinding activity across three independent experiments (Fig. 3D and E, and Supplementary Fig. S9). No detectable unwinding was observed for R493H in the single experiment performed, this result is therefore presented as qualitative evidence (Supplementary Fig. S9). Given that DHX30H562R is AT-Pase-deficient and DHX30R493H is impaired in RNA binding (20,21) these results show that DHX30-mediated R-loop unwinding requires both ATP hydrolysis and RNA substrate engagement, and that disease-associated *DHX30* variants are functionally defective in this activity.

Together, these data demonstrate that DHX30 is a bona fide R-loop helicase with defined structural and polarity requirements, providing a direct mechanistic explanation for the accumulation of RNA:DNA hybrids observed upon DHX30 deficiency. Given the intrinsic asymmetry of R-loops and models predicting exposure of the RNA 3′ terminus during transcription-replication conflicts (44), these biochemical properties are compatible with a role for DHX30 at R-loop structures formed under replication stress. Furthermore, impaired unwinding by the disease-associated variants suggests that defective R-loop processing contributes to DHX30-associated pathologies.

### DHX30 associates with a nuclear protein network linked to R-loop biology, including DHX9

R-loop metabolism is orchestrated by multicomponent assemblies of RNA-binding proteins, RNA helicases, and DNA repair factors, many of which are recruited to sites of transcription-replication conflicts (45,46). We therefore reasoned that DHX30 might function within a broader R-loop-associated protein network. To characterize the nuclear DHX30-associated protein network, DHX30-GFP was immunoprecipitated from nuclear fractions of HEK293T cells expressing the fusion protein, followed by mass spectrometry analysis. After filtering against control pulldowns, we identified 134 high-confidence DHX30-associated proteins (Supplementary Table 4). Proteins were retained based on detection of at least four unique peptides and a CHIMERYS identification score ≥25. These criteria support confident protein identification but do not distinguish direct binding from association within larger complexes. Functional enrichment analysis of the nuclear protein network associated with DHX30 revealed a strong overrepresentation of pathways related to RNA metabolism and translation, as identified by both Reactome pathway analysis and Gene Ontology enrichment. In addition, pathways associated with developmental processes and cellular stress responses were significantly enriched, together with a broad representation of proteins involved in gene expression regulation (Supplementary Fig. S10).

To further evaluate the relationship between DHX30 and R-loop biology, we cross-referenced the DHX30 nuclear protein network with the curated “R-loop regulators” dataset from the R-loop Base project (47). This analysis revealed that ∼64%, 87 of the 134 DHX30 interactors overlapped with proteins previously implicated in R-loop biology. Application of a more stringent confidence threshold identified a core subset of ten high-confidence proteins associated with R-loop biology, including multiple RNA helicases (DHX9, DDX1, DDX17, DDX18 and DDX21), the transcription-associated topoisomerase TOP1, nucleolin (NCL), the spliceosomal component SNRPA1, the chromatin-associated factor SAFB2 and the replication clamp PCNA (Fig. 4A).

**Fig. 4.**
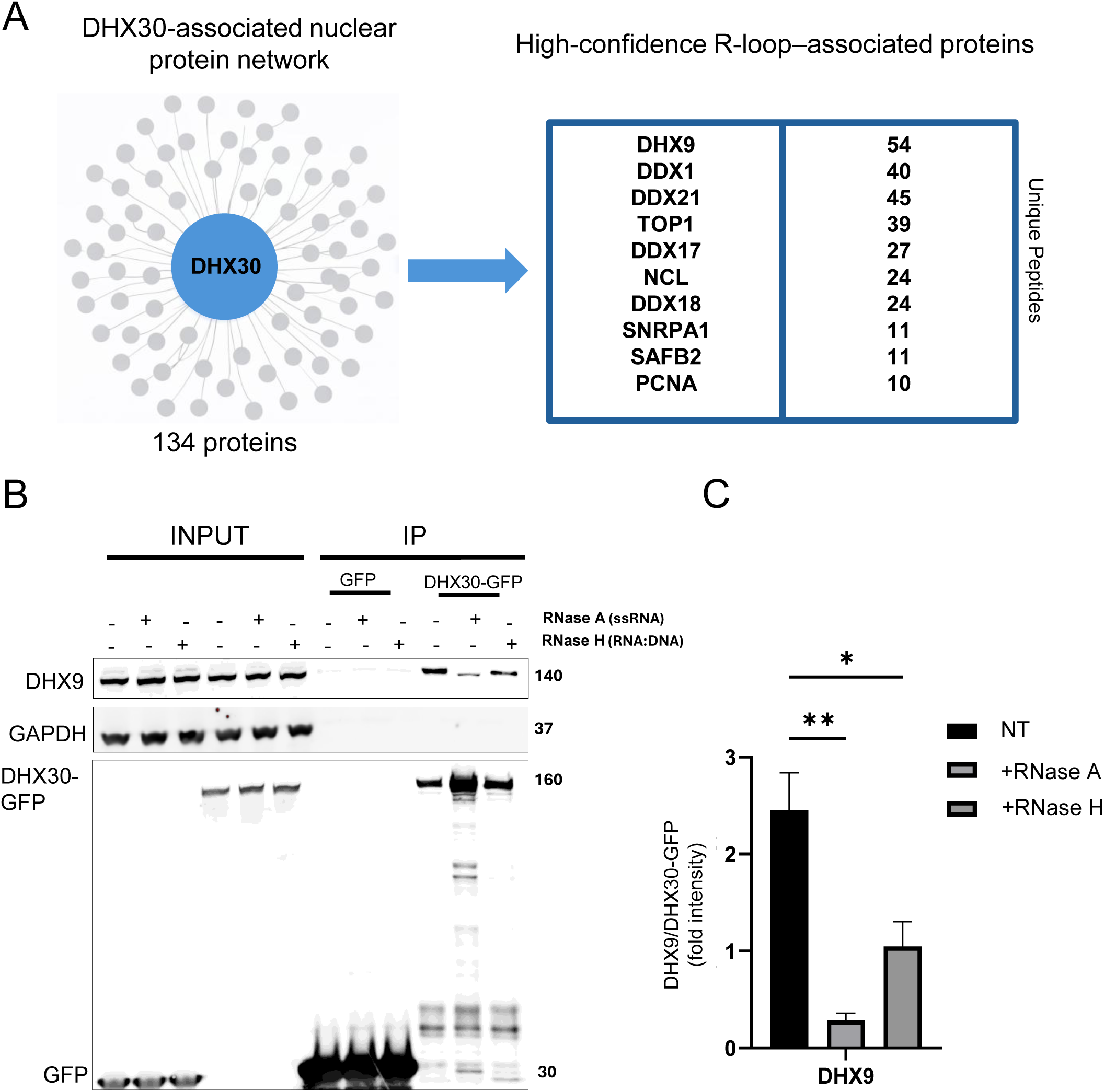
Mass spectrometry analysis of the DHX30-associated nuclear protein network. (**A**) Schematic representation of the proteomic analysis of the DHX30-associated nuclear proteins in HEK293T cells. DHX30 is shown as the central node, with DHX30-associated proteins depicted as surrounding nodes (134 proteins). Table listing high-confidence R-loop-associated proteins identified among DHX30 co-purifying proteins. Proteins are ranked based on the number of unique peptides detected, as indicated on the right. (**B**) Co-immunoprecipitation analysis of DHX9 co-purification with DHX30-GFP. Whole-cell lysates from HEK293T cells expressing DHX30-GFP or GFP alone were treated *in vitro* with either RNase A (in high salt conditions for the degradation of single-stranded RNA) or RNase H (for the degradation of RNA:DNA hybrids) prior to immunoprecipitation. (**C**) Quantification of DHX9 co-immunoprecipitation with DHX30-GFP under untreated (NT), RNase A treated, and RNase H treated conditions. Co-immunoprecipitated DHX9 signals were normalized to the corresponding recovered DHX30-GFP signals. Bars represent mean ± SEM from 3 independent experiments. Statistical analysis was assessed using one-way ANOVA followed by Dunnett’s multiple comparison test (*P < 0.05; **P < 0.01).

Among these candidates, we focused on DHX9, which had the highest number of unique peptides among the highlighted R-loop-associated proteins and shares significant sequence and functional homology with DHX30 (12). Notably, monoallelic missense variants in *DHX9* have been linked to neurodevelopmental disorders with phenotypic overlap to *DHX30*-associated conditions, suggesting potential functional convergence between these two helicases (48). To validate the association between DHX30 and DHX9, we performed co-immunoprecipitation assays using whole-cell lysates from cells expressing DHX30-GFP or GFP only. Immunoprecipitation of DHX30-GFP resulted in robust coprecipitation of endogenous DHX9 (Fig. 4B). Affinity purification of DHX30-GFP from nuclear extracts further confirmed that the DHX30–DHX9 interaction occurs specifically within the nuclear compartment (Supplementary Fig. S11a). Consistently, reciprocal co-immunoprecipitation experiments demonstrated that pulldown of DHX9-GFP efficiently recovered endogenous DHX30, supporting the specificity of this interaction (Supplementary Fig. S11b).

To delineate the regions of DHX30 required for DHX9 association, we performed domain-mapping experiments using a series of DHX30 truncation constructs. Co-immunoprecipitation assays revealed that DHX9 binding depends on the N-terminal RNA-binding domain of DHX30 (Double-stranded RNA-binding domains; dsRBD), suggesting a role for RNA in stabilizing this interaction (Supplementary Fig. S11c). Indeed, several previous studies suggested that DHX30 associated with proteins in an RNA dependent manner (49,50). Thus, to assess whether the DHX30-DHX9 association is mediated by RNA or RNA:DNA hybrids, cell lysates were treated *in vitro* with RNase A, which degrades single-stranded RNA, or RNase H, which selectively degrades RNA within DNA-RNA hybrids, prior to immunoprecipitation. Notably, treatment with either RNase A or RNase H significantly reduced the interaction (Fig. 4C), indicating that RNA species, including DNA-RNA hybrids, contribute to DHX30-DHX9 complex formation.

Furthermore, to assess whether RNA contributes to the association of DHX30 with other proteins, whole-cell lysates from HEK293T cells expressing DHX30-GFP were treated with RNase A prior to GFP-Trap immunoprecipitation. Coomassie staining indeed revealed markedly reduced recovery of co-purifying proteins following this treatment (Supplementary Fig. S12). Thus, these findings suggest that RNA broadly contributes to the formation of DHX30-associated protein complexes. Together, these data place DHX30 within a nuclear protein network enriched for proteins linked to RNA and R-loop biology and identify DHX9 as a prominent DHX30 associated factor.

### DHX30 suppresses R-loop-associated DNA damage and maintains genome stability

R-loops are a major source of transcription-replication conflicts, which can induce replication stress and genomic instability (51). Conversely, DNA damage and replication stress can further promote R-loop accumulation, thereby establishing a self-reinforcing cycle of genome instability (8). To determine whether RNA:DNA hybrid accumulation upon DHX30-deficiency is associated with increased DNA damage, we quantified two well-established markers of DNA damage: γH2AX, the phosphorylated form of histone H2AX (Ser139), which marks sites of single- and double-strand DNA breaks (52), and 53BP1, a key DNA damage response factor that accumulates in discrete nuclear foci at double-strand breaks (53). Immunofluorescence analysis of U2OS cells transiently depleted of DHX30 by siRNA revealed a significant increase in nuclear γH2AX signal intensity and a higher proportion of 53BP1-positive nuclear foci compared with siControl-treated cells (Fig. 5A-C). Notably, this increase was detectable under basal conditions and was further exacerbated following treatment with replication stress-inducing agents, APH and CPT (Fig. 5A-C), consistent with increased formation of replication-associated DNA double-strand breaks.

**Fig. 5.**
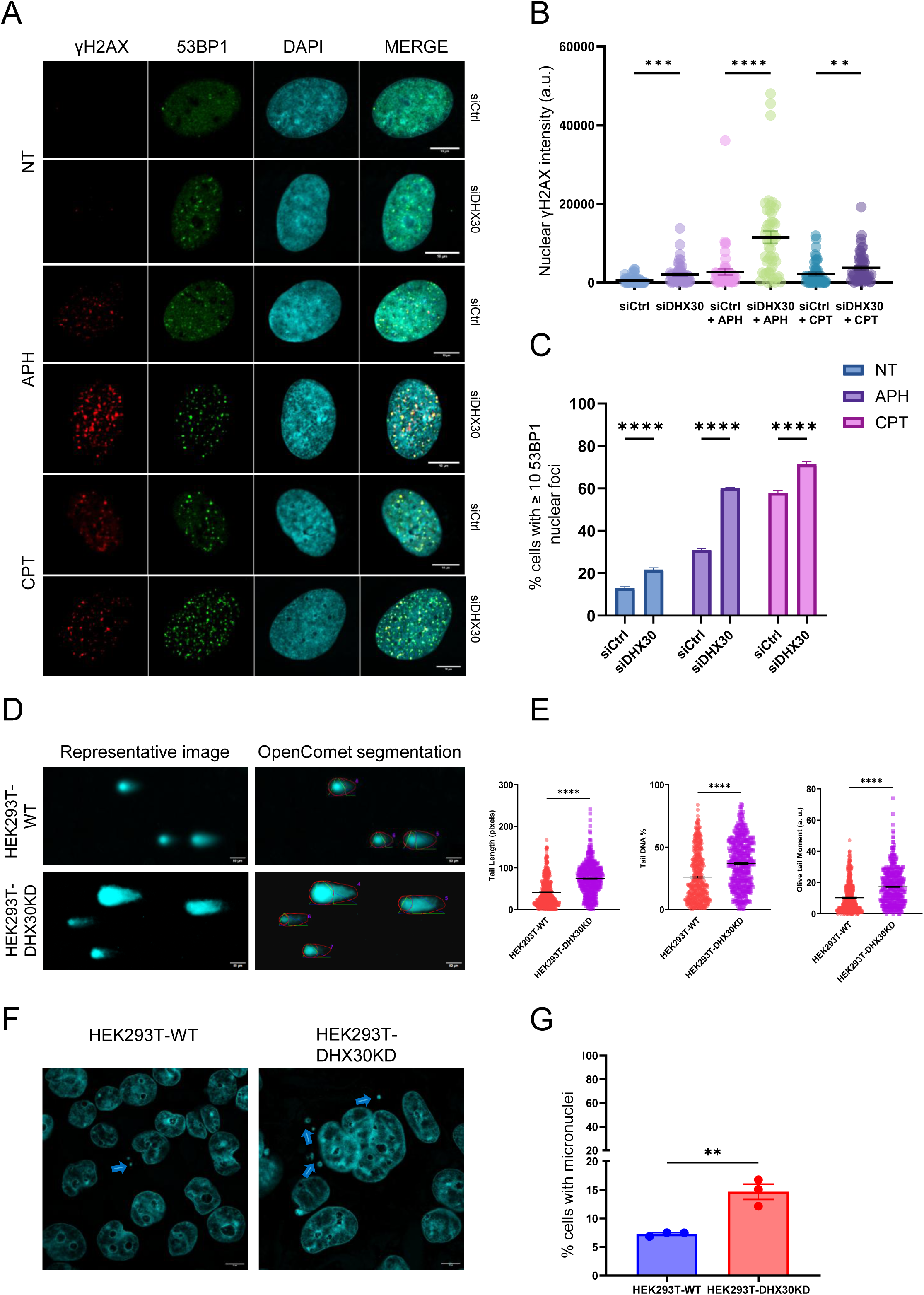
DHX30 deficiency causes DNA damage and genome instability. **(A)** Representative single-plane immunofluorescence images of U2OS cells transfected with control siRNA (siCtrl) or DHX30-targeting siRNA (siDHX30) under untreated (NT) conditions or following treatment with aphidicolin (APH) or camptothecin (CPT). Cells were stained for γH2AX (red), 53BP1 (green), and DAPI (cyan). Merged images are shown in the right panels. Scale bars, 10 μm. **(B)** Quantification of nuclear γH2AX fluorescence intensity expressed as arbitrary units (a.u.) in cells treated as indicated. γH2AX intensity was quantified using Fiji. Each point represents one cell. Approximately 50 cells per condition, pooled from three independent biological experiments, were analyzed. Statistical significance was assessed using the Kruskal-Wallis test followed by Dunn’s multiple-comparisons test for the indicated pairwise comparisons (**P < 0.01; ***P < 0.001; ****P < 0.0001). **(C)** Percentage of nuclei containing ≥10 53BP1 foci in siCtrl and siDHX30 cells under the indicated conditions. Data are presented as mean ± SEM from three independent biological experiments. Statistical significance was assessed using ordinary two-way ANOVA followed by Šídák’s multiple-comparisons test, comparing siCtrl and siDHX30 within each treatment condition (****P < 0.0001). **(D)** Representative images from neutral comet assays performed in HEK293T-WT and HEK293T-DHX30KD cells, together with the corresponding OpenComet segmentation used for analysis. Scale bars, 50 μm. **(E)** Quantification of tail length, Tail DNA (%), and Olive tail moment using OpenComet. Each dot represents an individual comet (number of comets=500); pooled data from four independent experiments are shown. Statistical significance was determined using a two-tailed Welch’s t-test (****P < 0.0001). **(F)** Representative fluorescence images of HEK293T-WT and HEK293T-DHX30KD cells stained with DAPI. Arrows indicate micronuclei. Scale bars, 10 μm. **(G)** Quantification of the percentage of cells containing micronuclei in HEK293T-WT and HEK293T-DHX30KD cells. Data are presented as mean ± SEM from three independent biological experiments. Statistical significance was assessed using an unpaired two-tailed t-test (**\*\***P < 0.01).

We next examined RPA32 phosphorylation at Ser4/Ser8, a marker of single-stranded DNA accumulation and DNA end resection (54). The percentage of cells containing >5 nuclear RPA32 pS4/S8 foci did not differ significantly between control and DHX30-deficient HEK293T cells (Supplementary Fig. S13). Thus, the increased γH2AX and 53BP1 signals were not accompanied by a detectable increase in this RPA32 phosphorylation readout.

To independently assess DNA double-strand break formation, we performed neutral comet assays in HEK293T cells, an established approach for detecting DNA double-strand breaks (55). DHX30-deficient cells showed significant increases in comet tail length, percentage of DNA in the tail, and Olive tail moment compared with controls, consistent with an increased burden of DNA double-strand breaks (Fig. 5D and E). To determine whether DHX30 deficiency causes broader genome instability, we quantified micronucleus formation, an established indicator of chromosomal damage (56). DHX30-deficient HEK293T cells showed a significant increase in the proportion of nuclei with associated micronuclei compared with controls (Fig. 5F and G), demonstrating that the consequences of DHX30 deficiency extend beyond DNA break accumulation to broader genome instability.

To determine whether DHX30 deficiency affects DNA synthesis, we performed 5-ethynyl-2’-deoxyuridine (EdU) pulse-labelling. DHX30-deficient cells exhibited a significant reduction in EdU incorporation compared with control cells (Supplementary Fig. S14a and b). We then asked whether DHX30 deficiency also increases DNA damage in cells undergoing DNA replication by quantifying 53BP1 foci in EdU-positive cells. This analysis revealed a marked increase in 53BP1 foci in EdU-positive DHX30-deficient cells compared with EdU-positive controls (Supplementary Fig. S14c). These findings demonstrate reduced DNA synthesis and increased DNA damage in actively replicating DHX30-deficient cells, consistent with impaired DNA replication (57).

Next, to directly test whether the DNA damage phenotype associated with DHX30 deficiency is cause by excessive RNA:DNA hybrid accumulation, we performed a series of rescue experiments in DHX30-deficient cells. Overexpression of nuclear RNase H1 in DHX30KD cells, which selectively degrades the RNA strand of RNA:DNA hybrids, resulted in a marked reduction of nuclear S9.6 signal and a concomitant decrease in 53BP1 foci, indicating that the DNA damage phenotype arises from excessive R-loop accumulation (Fig. 6B and C). These findings are consistent with the established role of RNase H in suppressing R-loop-induced genome instability (58). Re-expression of DHX30 in DHX30-deficient cells similarly reduced nuclear S9.6 signal and 53BP1 foci and increased EdU positivity (Fig. 6; Supplementary Fig. S14d and e), further supporting a direct link between reduced DHX30, RNA:DNA accumulation, DNA damage and DNA synthesis. Interestingly, overexpression of DHX9 in DHX30KD cells, a well-characterized RNA helicase involved in R-loop homeostasis (19), also significantly reduced both nuclear S9.6 signal and 53BP1 foci, suggesting a partial compensatory capacity between these helicases (Fig. 6B and C).

**Fig. 6.**
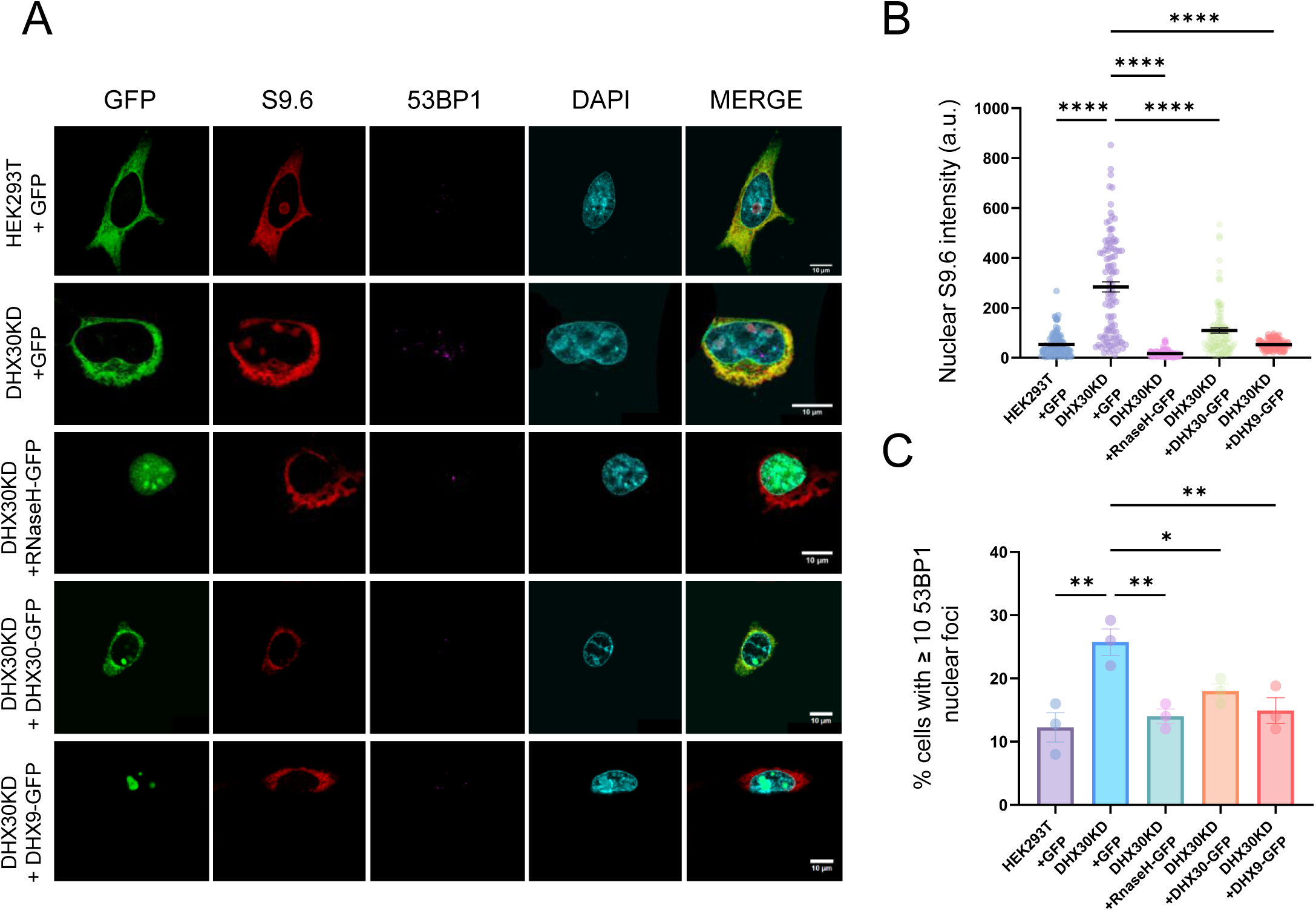
Overexpression of RNase H1, DHX30 and DHX9 rescue S9.6-positive nuclear signal and DNA damage in DHX30-deficient cells. (**A**) Representative single-plane immunofluorescence images showing DAPI (cyan), GFP (green), 53BP1 (magenta) and S9.6 (red) staining in HEK293T wild-type and DHX30 knockdown (DHX30KD) cells transfected with GFP only, RNase H1-GFP, DHX30-GFP or DHX9-GFP, as indicated. Merged images are shown in the right panels. Scale bars, 10 μm. (**B**) Quantification of nuclear S9.6 fluorescence intensity expressed as arbitrary units (a.u.) in cells treated as in (**A**). S9.6 intensity was quantified using Fiji. Each dot represents one nucleus. Approximately 100 cells per condition, pooled from three independent biological experiments, were analyzed. Statistical significance was assessed using ordinary one-way ANOVA followed by Dunnett’s multiple-comparisons test, comparing each condition with DHX30KD + GFP (****P < 0.0001). (**C**) Counting of the percentage of nuclei containing ≥ 10 53BP1 foci in the indicated conditions. Bars represent mean ± SEM from three biologically independent experiments. Statistical significance was assessed using one-way ANOVA followed by Dunnett’s multiple comparison test, comparing each condition with DHX30KD + GFP (*P < 0.05; **P < 0.01).

Finally, we asked whether the disease-associated *DHX30* variants R493H and H562R, which were defective in R-loop unwinding *in vitro*, could restore genome maintenance in DHX30-deficient cells. Expression of either variant failed to significantly reduce the proportion of cells with elevated 53BP1 foci compared with the GFP control, indicating that neither variant effectively rescued the DNA damage phenotype (Supplementary Fig. S15a). We next assessed whether these variants could restore DNA synthesis. Neither R493H nor H562R significantly increased the proportion of EdU-positive cells relative to the GFP control (Supplementary Fig. S15b). Expression of the wild-type and variant constructs was confirmed by immunoblotting (Supplementary Fig. S15c). Together, these findings link the impaired R-loop unwinding activity of disease-associated *DHX30* variants to a failure to rescue the cellular consequences of DHX30 deficiency.

Collectively, these results support a model in which DHX30 limits R-loop-induced replication stress and consequent genome instability, and suggest that DHX30 and DHX9 share overlapping functions in the maintenance of R-loop homeostasis and genome stability. The failure of the disease-associated R493H and H562R variants to rescue DNA damage and DNA synthesis defects further suggests that these protective functions are compromised in *DHX30*-associated neurodevelopmental disorder.

### DHX30 deficiency disrupts transcriptional homeostasis and induces a stress-associated gene expression program

Given the effects of DHX30 deficiency on RNA:DNA hybrid homeostasis and genome stability, we next examined whether downregulation of DHX30 impacts global transcriptional homeostasis. To this end, we performed RNA-sequencing (RNA-seq) analyses in control and DHX30-deficient cells. Differential expression analysis revealed significant changes in steady-state RNA levels upon DHX30 deficiency, with 754 genes significantly upregulated and 714 genes downregulated relative to control cells (FDR < 0.001 & absolute logFC > 1) (Fig. 7A and Supplementary Table 5). Notably, we observed a 50% reduction of DHX30 on the mRNA level in DHX30 deficient cells (Supplementary Fig. S16 and Supplementary Table 5). Gene Ontology enrichment analysis of the deregulated gene set identified multiple biological processes related to neuronal development and function, including axonogenesis, neuron projection guidance, axon guidance, and postsynaptic organization (Fig. 7B). To determine which processes were associated with increased or decreased gene expression, upregulated and downregulated genes were analysed separately. Downregulated genes were enriched for neuronal and synaptic processes, including axonogenesis, neuron projection guidance, postsynaptic organization, and regulation of membrane potential, whereas upregulated genes were predominantly associated with broader developmental and morphogenetic processes (Supplementary Fig. S17a and b). The enrichment of neuronal and synaptic processes among downregulated genes is notable in the context of DHX30-associated neurodevelopmental disorder (20,21).

**Fig. 7.**
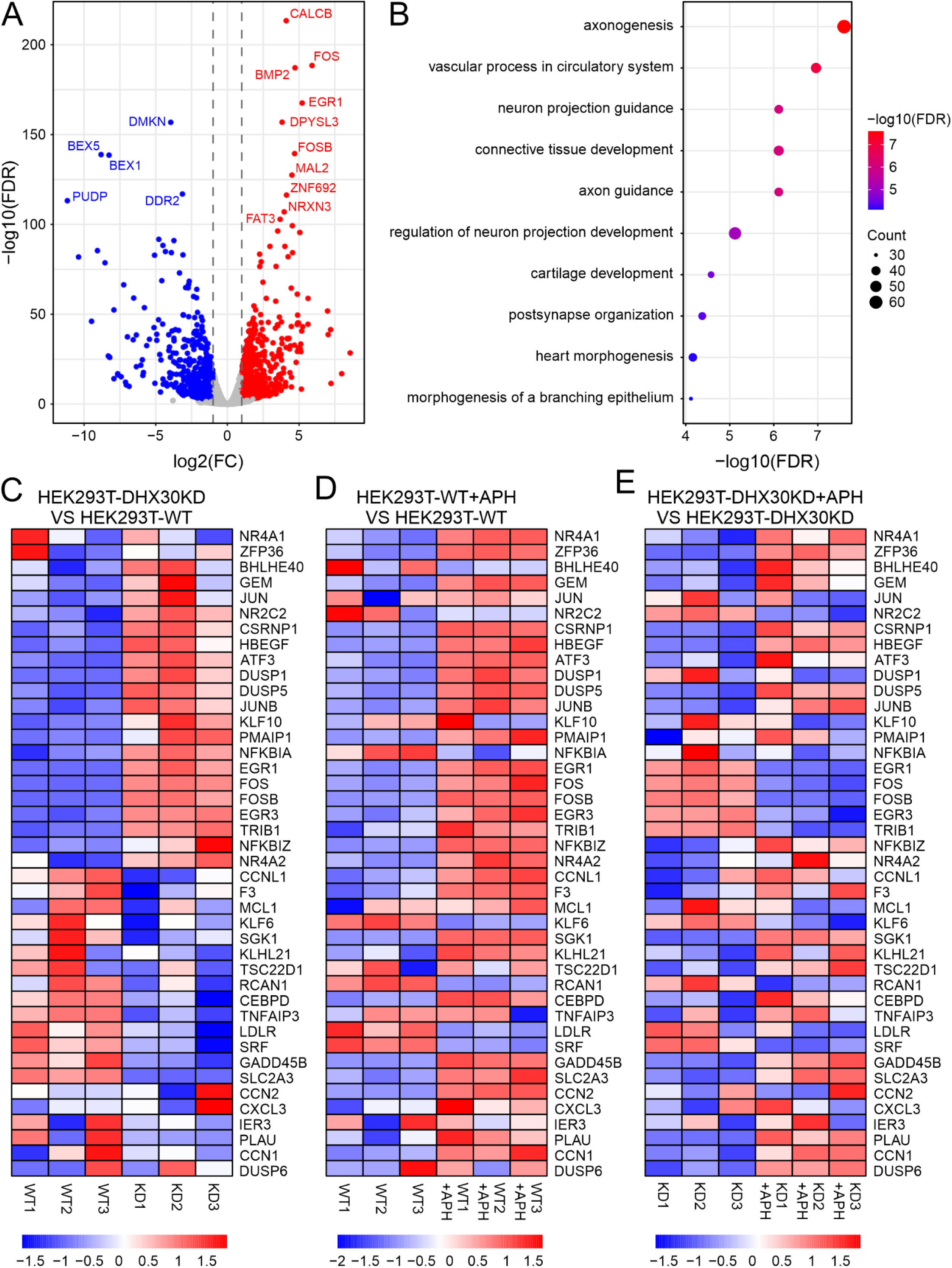
DHX30 deficient cells show significantly dysregulated transcriptome. (**A**) Volcano plot showing differential gene expression in HEK293T-DHX30 knockdown (DHX30KD) cells compared with HEK293T wild-type cells. Significantly upregulated and downregulated genes are shown in red and blue, respectively (FDR < 0.001 & absolute logFC > 1). The top genes are labelled. **(B)** Top 10 pathways from Gene Ontology (GO) enrichment analysis of significantly deregulated genes in DHX30KD cells. Dot size indicates the number of genes associated with each term, and colour represents statistical significance (−log10 FDR). (**C**) Heatmap showing expression levels of selected immediate-early stress-responsive genes in HEK293T-DHX30KD versus WT cells. Individual biological replicates are shown. (**D**) Heat-map of selected immediate-early stress-responsive genes expression changes in HEK293TWT cells treated with aphidicolin (APH) compared with untreated WT cells. (**E**) Heatmap showing selected immediate-early stress-responsive genes expression changes in DHX30KD cells upon APH treatment compared with untreated DHX30KD cells. Colour scale represents relative gene expression levels, with red indicating higher expression and blue indicating lower expression.

Among the most strongly upregulated transcripts, we observed *FOS*, *EGR1* and *FOSB* (Fig. 7A), members of the immediate-early gene (IEG) family that are rapidly and transiently induced in response to cellular stress or proliferative signals (59). This prompted us to specifically assess the expression of IEGs using a previously curated gene set (60). Consistent with a broader activation of this program, several additional IEGs including *EGR3*, *DUSP1*, *DUSP5*, *ATF3* and *PMAIP1* (NOXA) were significantly upregulated (logFC > 1) in DHX30-depleted cells as compared to controls (Fig. 7C and Supplementary Table 5). RT-qPCR analysis of two representative genes, *EGR1* and *PMAIP1*, confirmed their increased transcript abundance in DHX30-deficient cells, independently supporting the RNA-seq findings (Supplementary Fig. S18).

Together, these transcriptional changes are consistent with activation of a stress-responsive gene program in DHX30-deficient cells. Persistent R-loops can form physical obstacles to replication forks during S phase and thereby promote replication stress (61). Replication stress, in turn, can elicit rapid stress-response transcriptional programs that include induction of immediate-early genes such as *FOS* (62).

We next examined how replication stress influences IEG expression dynamics in the context of DHX30 deficiency. Control cells treated with APH exhibited a robust and coordinated transcriptional response, characterized by strong induction of immediate-early and stress-adaptive genes, including *ATF3*, *EGR1*, *FOS*, *FOSB*, *DUSP1*, *DUSP5*, *JUN*, and *JUNB* (Fig. 7D). In contrast, DHX30-deficient cells displayed an altered transcriptional response, with attenuated or heterogeneous induction of IEGs following APH treatment. Notably, several IEG transcripts that were already elevated under basal conditions in DHX30-deficient cells, particularly *EGR1*, *FOS*, *FOSB*, *EGR3* and *TRIB1* were downregulated upon replication stress (Fig. 7E). These data indicate that DHX30 deficiency is associated with inappropriate basal activation of stress-responsive genes and impaired induction of replication stress-adaptive gene expression programs.

Collectively, these findings reveal extensive changes in steady-state gene expression upon DHX30 deficiency that converges on pathways involved in neuronal development, cell signalling, and stress-responsive gene regulation, and demonstrate that DHX30 deficiency disrupts both basal and replication stress-induced transcriptional programs, consistent with impaired R-loop resolution during DNA replication.

## DISCUSSION

In this study, we identify the DExH-box RNA helicase DHX30 as a nuclear regulator of R-loop homeostasis, particularly under conditions of replication stress. We show that, beyond its previously described cytoplasmic and mitochondrial roles, DHX30 localizes to the nucleus, where it overlaps with S9.6-positive regions, and limits the accumulation of RNA:DNA hybrids in both human cells and zebrafish. DHX30 deficiency leads to RNA:DNA hybrid accumulation, replication-associated DNA double-strand breaks, micronucleus formation, impaired DNA synthesis and maladaptive transcriptional responses, suggesting that DHX30 contributes to cellular mechanisms that mitigate transcription-replication conflicts.

Mechanistically, our biochemical analyses demonstrate that DHX30 exhibits substrate selectivity, as it directly unwinds R-loop substrates carrying a 3’single-stranded RNA overhang in an ATP-dependent manner (Fig. 3). This identifies DHX30 as a bona fide R-loop helicase with polarity requirements, consistent with the loading and translocation properties of DExH-box helicases (63–65). These findings provide a possible molecular explanation for the function of DHX30 at transcription-replication conflict sites, where models predict exposure of RNA 3′termini (66–68). Consistently, DHX30 shows preferential nuclear enrichment following aphidicolin or camptothecin treatment, conditions that induce replication stress and promote transcription-replication conflicts (Fig. 1C). The mechanism underlying this nuclear accumulation however still remains unknown and may involve changes in nuclear import, export, retention or protein stability (69–71). Furthermore, DHX30 deficiency induces RNA:DNA hybrid accumulation in cellular and animal models (Fig. 2). This behaviour is consistent with that of several RNA helicases implicated in R-loop control, including DDX17, DDX18, DDX3X, DDX21, and DHX9, whose loss similarly leads to RNA:DNA hybrid accumulation and genome instability (19,72–76). Our proteomic and functional data further place DHX30 within a nuclear RNA and R-loop regulatory network and suggest overlapping functions among RNA helicases in mitigating R-loop-associated threats to genome integrity. These findings are consistent with previous proteomic studies that identified DHX30 as an R-loop-associated protein (40)(77). In line with previous studies (49,50), the RNase A sensitivity of DHX30-associated complexes further suggests that RNA contributes substantially to these assemblies.

Consistent with its role in limiting R-loop accumulation DHX30 deficiency leads to pronounced replication-associated genome instability. DHX30-deficient cells exhibit impaired DNA synthesis, as evidenced by reduced EdU incorporation, together with elevated DNA damage signalling, manifested by increased nuclear γH2AX intensity and 53BP1 nuclear foci under basal conditions and following APH and CPT treatment (Fig. 5A-C and Supplementary Fig. S14). The increased burden of 53BP1 foci in EdU-positive DHX30-deficient cells further supports an association between impaired DNA replication and DNA damage. Importantly, neutral comet assays independently demonstrated increased DNA fragmentation consistent with an elevated burden of double-strand breaks, while increased micronucleus formation revealed broader chromosomal damage (Fig. 5D–G). These features are consistent with a model of replication fork instability and double-strand break formation arising from unresolved transcription-replication conflicts (78). Importantly, 53BP1 accumulation is efficiently suppressed by RNase H1 expression or by reintroduction of wild-type DHX30, supporting a contribution of excessive R-loops to DNA damage in the absence of DHX30 (Fig. 6). Both RNase H1 and wild-type DHX30 also increased EdU positivity, linking hybrid homeostasis to the DNA synthesis phenotype (Supplementary Fig. S14d and e). Partial rescue by DHX9 overexpression further supports functional overlap among nuclear RNA helicases in mitigating R-loop related replication stress. These complementation results place DHX30 function within the broader context of established RNA:DNA hybrid-processing pathways. RNase H1 and RNase H2 remove hybrids by degrading their RNA component, whereas RNA helicases can promote their resolution through strand separation (16,79). Notably, pathogenic variants in genes encoding RNase H2 subunits cause a neuroinflammatory, Aicardi-Goutières syndrome, and RNase H2 deficiency leads to co-transcriptional R-loop accumulation, impaired gene expression and DNA damage (80). Among disease-associated RNA helicases (12), an NDD-associated *DHX9* variant increases R-loop accumulation and DNA double-strand breaks (48), while DDX3X processes R-loop substrates and stimulates RNase H2 activity (76). Together, these observations raise the possibility that disturbed R-loop homeostasis may contribute to the cellular consequences of dysfunction across several neurologically relevant RNA-processing factors.

In parallel with replication-associated genome instability, DHX30 deficiency profoundly alters the steady-state expression of immediate-early genes (IEGs). Separate Gene Ontology analyses additionally revealed enrichment of neuronal and synaptic processes among down-regulated genes, whereas upregulated genes were associated with broader developmental and morphogenetic processes (Supplementary Fig. S17). Although obtained in HEK293T cells, these findings are notable in the context of DHX30-associated neurodevelopmental disorder. DHX30KD cells exhibit inappropriate basal activation of stress-responsive IEGs, including *FOS*, *FOSB*, and *EGR1*, even in the absence of exogenous replication perturbation (Fig. 7C). These expression abnormalities may be particularly detrimental in rapidly proliferating or differentiating neural progenitors (81). Given the concomitant accumulation of R-loops and DNA damage under these conditions, this basal IEG activation is consistent with chronic replication-associated stress signalling that may be driven, at least in part, by unresolved RNA:DNA hybrids (82), and is in line with evidence that replication stress can elicit immediate-early transcriptional responses (83). Notably, upon aphidicolin-induced replication challenge, DHX30-deficient cells fail to mount a coordinated stress-adaptive transcriptional response and instead display attenuation or downregulation of IEG expression (Fig. 7E). This impaired response may reflect a state of chronic replication stress in DHX30-deficient cells, in which persistent R-loop accumulation preconditions the transcriptional stress response and limits the capacity of cells to dynamically activate immediate-early gene programs upon additional replication perturbation. Our findings support a model in which DHX30-mediated control of R-loop accumulation contributes to the coordination of genome stability with stress-responsive transcriptional programs.

Moreover, our biochemical assays show that disease-associated *DHX30* missense variants are defective in R-loop unwinding (Fig. 3D and E, and Supplementary Fig. S9), directly linking pathogenic variants to impaired RNA:DNA hybrid resolution. This defect is consistent with prior studies showing that these variants are unable to unwind canonical RNA duplexes, reflecting a fundamental loss of helicase activity (20). Furthermore, in the cellular complementation experiments, R493H and H562R did not significantly reduce 53BP1 foci or improve EdU positivity relative to the GFP control (Supplementary Fig. S15). The failure of these variants to rescue these cellular phenotypes may reflect both their impaired R-loop unwinding activity and their previously reported sequestration in cytoplasmic stress granules (20,21), which would be expected to limit their nuclear availability. Thus, defective catalytic activity and altered subcellular localization may jointly compromise the ability of these variants to maintain nuclear R-loop homeostasis. However, the precise contribution of these defects to the human neurodevelopmental phenotype remains to be fully defined. To address this question, we are currently establishing patient-derived induced pluripotent stem cells and differentiated neurons, which will enable analysis of DHX30-dependent R-loop regulation, transcriptional control, and genome stability in disease-relevant neural contexts. Comprehensive investigation of these patient-derived neuronal models will require substantial experimental effort and will constitute a central focus of our future work.

In summary, our study identifies DHX30 as a conserved regulator of RNA:DNA homeostasis that directly unwinds R-loop substrates, safeguards genome stability and coordinates stress-responsive transcription. Our findings place DHX30 within a network of RNA helicases that resolve R-loop-mediated transcription-replication conflicts and thereby maintain genomic integrity. The impaired R-loop unwinding activity and failure of disease-associated variants to rescue cellular defects provide a mechanistic framework for investigating how disrupted nuclear R-loop regulation may contribute to *DHX30*-associated neurodevelopmental disorder.

## Data availability

The RNA sequencing data have been deposited in GEO under accession number GSE322799.

---

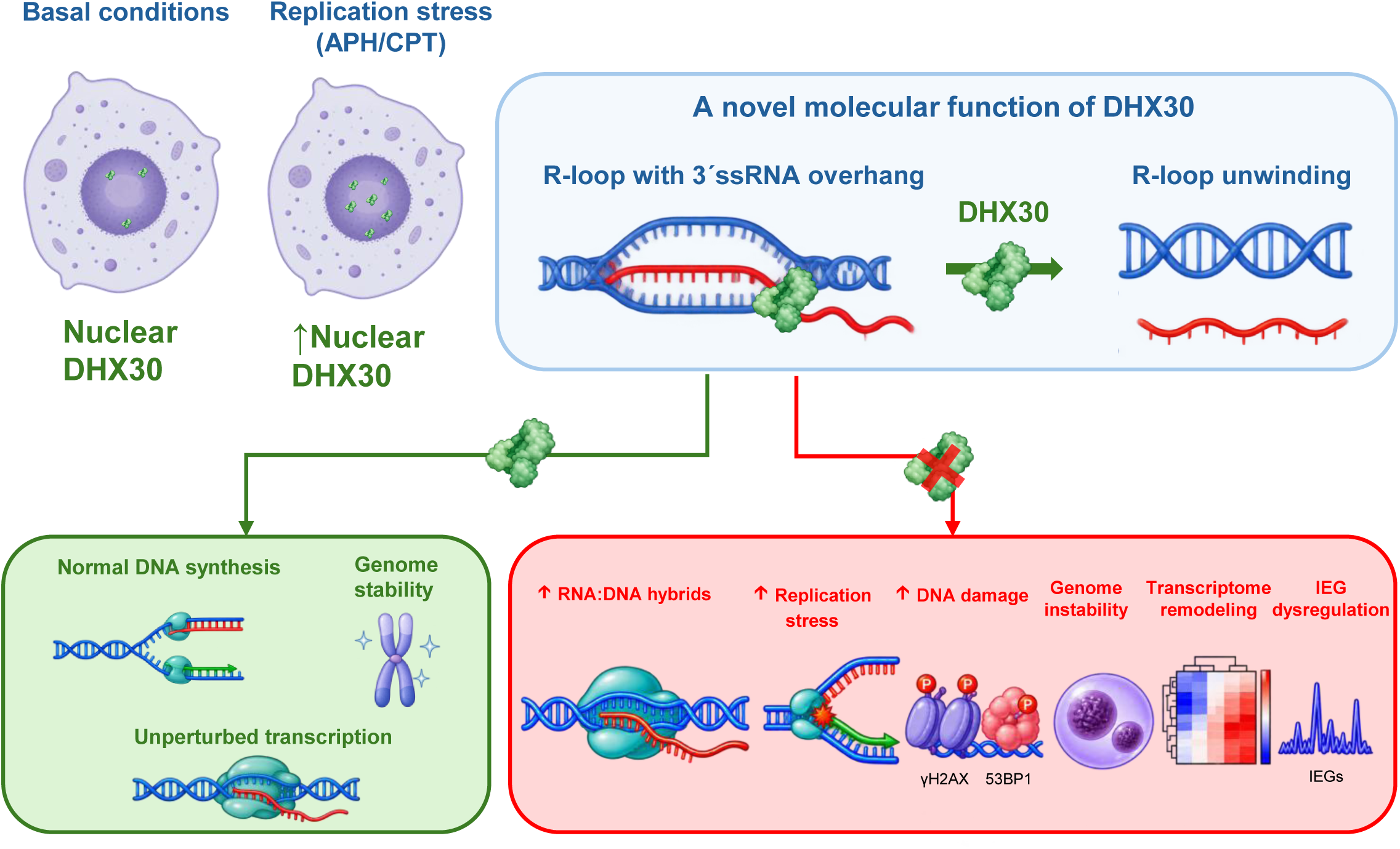

## References

1. Gorbalenya, A.E. and Koonin, E.V. (1993) Helicases - Amino-Acid-Sequence Comparisons and Structure-Function-Relationships. Curr Opin Struc Biol, 3, 419–429.

2. Singleton, M.R., Dillingham, M.S. and Wigley, D.B. (2007) Structure and mechanism of helicases and nucleic acid translocases. Annu Rev Biochem, 76, 23–50.

3. Linder, P. and Jankowsky, E. (2011) From unwinding to clamping - the DEAD box RNA helicase family. Nat Rev Mol Cell Biol, 12, 505–516.

4. Linder, P., Lasko, P.F., Ashburner, M., Leroy, P., Nielsen, P.J., Nishi, K., Schnier, J. and Slonimski, P.P. (1989) Birth of the D-E-a-D Box. Nature, 337, 121–122.

5. Jankowsky, E., Gross, C.H., Shuman, S. and Pyle, A.M. (2001) Active disruption of an RNA-protein interaction by a DExH/D RNA helicase. Science, 291, 121–125.

6. Bourgeois, C.F., Mortreux, F. and Auboeuf, D. (2016) The multiple functions of RNA helicases as drivers and regulators of gene expression. Nat Rev Mol Cell Biol, 17, 426–438.

7. Jarmoskaite, I. and Russell, R. (2014) RNA helicase proteins as chaperones and remodelers. Annu Rev Biochem, 83, 697–725.

8. Santos-Pereira, J.M. and Aguilera, A. (2015) R loops: new modulators of genome dynamics and function. Nat Rev Genet, 16, 583–597.

9. Sanz, L.A., Hartono, S.R., Lim, Y.W., Steyaert, S., Rajpurkar, A., Ginno, P.A., Xu, X. and Chedin, F. (2016) Prevalent, Dynamic, and Conserved R-Loop Structures Associate with Specific Epigenomic Signatures in Mammals. Mol Cell, 63, 167–178.

10. Aguilera, A. and Garcia-Muse, T. (2012) R loops: from transcription byproducts to threats to genome stability. Mol Cell, 46, 115–124.

11. Huertas, P. and Aguilera, A. (2003) Cotranscriptionally formed DNA:RNA hybrids mediate transcription elongation impairment and transcription-associated recombination. Mol Cell, 12, 711–721.

12. Lederbauer, J., Das, S., Piton, A., Lessel, D. and Kreienkamp, H.J. (2024) The role of DEAD- and DExH-box RNA helicases in neurodevelopmental disorders. Front Mol Neurosci, 17, 1414949.

13. Bhattacharjee, R., Jolly, L.A., Corbett, M.A., Wee, I.C., Rao, S.R., Gardner, A.E., Ritchie, T., van Hugte, E.J.H., Ciptasari, U., Piltz, S., et al. (2024) Compromised transcription-mRNA export factor THOC2 causes R-loop accumulation, DNA damage and adverse neurodevelopment. Nat Commun, 15, 1210.

14. Yan, P., Liu, Z., Song, M., Wu, Z., Xu, W., Li, K., Ji, Q., Wang, S., Liu, X., Yan, K. et al. (2020) Genome-wide R-loop Landscapes during Cell Differentiation and Reprogramming. Cell Rep, 32, 107870.

15. Garcia-Muse, T. and Aguilera, A. (2019) R Loops: From Physiological to Pathological Roles. Cell, 179, 604–618.

16. Yang, S., Winstone, L., Mondal, S. and Wu, Y. (2023) Helicases in R-loop Formation and Resolution. J Biol Chem, 299, 105307.

17. Polenkowski, M., Allister, A.B., Burbano de Lara, S., Pierce, A., Geary, B., El Bounkari, O., Wiehlmann, L., Hoffmann, A., Whetton, A.D., Tamura, T., et al. (2023) THOC5 complexes with DDX5, DDX17, and CDK12 to regulate R loop structures and transcription elongation rate. iScience, 26, 105784.

18. Li, L., Germain, D.R., Poon, H.Y., Hildebrandt, M.R., Monckton, E.A., McDonald, D., Hendzel, M.J. and Godbout, R. (2016) DEAD Box 1 Facilitates Removal of RNA and Homologous Recombination at DNA Double-Strand Breaks. Mol Cell Biol, 36, 2794–2810.

19. Chakraborty, P. and Grosse, F. (2011) Human DHX9 helicase preferentially unwinds RNA-containing displacement loops (R-loops) and G-quadruplexes. DNA Repair (Amst), 10, 654–665.

20. Mannucci, I., Dang, N.D.P., Huber, H., Murry, J.B., Abramson, J., Althoff, T., Banka, S., Baynam, G., Bearden, D., Beleza-Meireles, A. et al. (2021) Genotype-phenotype correlations and novel molecular insights into the DHX30-associated neurodevelopmental disorders. Genome Med, 13, 90.

21. Lessel, D., Schob, C., Kury, S., Reijnders, M.R.F., Harel, T., Eldomery, M.K., Coban-Akdemir, Z., Denecke, J., Edvardson, S., Colin, E. et al. (2017) De Novo Missense Mutations in DHX30 Impair Global Translation and Cause a Neurodevelopmental Disorder. Am J Hum Genet, 101, 716–724.

22. Leone, S., Bar, D., Slabber, C.F., Dalcher, D. and Santoro, R. (2017) The RNA helicase DHX9 establishes nucleolar heterochromatin, and this activity is required for embryonic stem cell differentiation. EMBO Rep, 18, 1248–1262.

23. Lessel, D., Vaz, B., Halder, S., Lockhart, P.J., Marinovic-Terzic, I., Lopez-Mosqueda, J., Philipp, M., Sim, J.C., Smith, K.R., Oehler, J. et al. (2014) Mutations in SPRTN cause early onset hepatocellular carcinoma, genomic instability and progeroid features. Nat Genet, 46, 1239–1244.

24. Zhang, M., Jiao, Z., Florez, Y.C., Lessel, I., Li, X., Yao, M.A., Weigl, M., Ozimek, M., Dogruluk, T., Chen, K., et al. (2025) miRISC inhibition causes mitotic defects and synergizes with genotoxic agents in cancers. BioRxiv.

25. Ewels, P., Magnusson, M., Lundin, S. and Kaller, M. (2016) MultiQC: summarize analysis results for multiple tools and samples in a single report. Bioinformatics, 32, 3047–3048.

26. Chen, S., Zhou, Y., Chen, Y. and Gu, J. (2018) fastp: an ultra-fast all-in-one FASTQ preprocessor. Bioinformatics, 34, i884–i890.

27. Mudge, J.M., Carbonell-Sala, S., Diekhans, M., Martinez, J.G., Hunt, T., Jungreis, I., Loveland, J.E., Arnan, C., Barnes, I., Bennett, R. et al. (2025) GENCODE 2025: reference gene annotation for human and mouse. Nucleic Acids Res, 53, D966–D975.

28. Dobin, A., Davis, C.A., Schlesinger, F., Drenkow, J., Zaleski, C., Jha, S., Batut, P., Chaisson, M. and Gingeras, T.R. (2013) STAR: ultrafast universal RNA-seq aligner. Bioinformatics, 29, 15–21.

29. Li, B. and Dewey, C.N. (2011) RSEM: accurate transcript quantification from RNA-Seq data with or without a reference genome. Bmc Bioinformatics, 12.

30. Durinck, S., Moreau, Y., Kasprzyk, A., Davis, S., De Moor, B., Brazma, A. and Huber, W. (2005) BioMart and Bioconductor: a powerful link between biological databases and microarray data analysis. Bioinformatics, 21, 3439–3440.

31. Chen, Y., Chen, L., Lun, A.T.L., Baldoni, P.L. and Smyth, G.K. (2025) edgeR v4: powerful differential analysis of sequencing data with expanded functionality and improved support for small counts and larger datasets. Nucleic Acids Res, 53.

32. Xu, S., Hu, E., Cai, Y., Xie, Z., Luo, X., Zhan, L., Tang, W., Wang, Q., Liu, B., Wang, R. et al. (2024) Using clusterProfiler to characterize multiomics data. Nat Protoc, 19, 3292–3320.

33. Antonicka, H. and Shoubridge, E.A. (2015) Mitochondrial RNA Granules Are Centers for Posttranscriptional RNA Processing and Ribosome Biogenesis. Cell Rep, 10, 920–932.

34. Wang, Y. and Bogenhagen, D.F. (2006) Human mitochondrial DNA nucleoids are linked to protein folding machinery and metabolic enzymes at the mitochondrial inner membrane. J Biol Chem, 281, 25791–25802.

35. Singatulina, A.S., Hamon, L., Sukhanova, M.V., Desforges, B., Joshi, V., Bouhss, A., Lavrik, O.I. and Pastre, D. (2019) PARP-1 Activation Directs FUS to DNA Damage Sites to Form PARG-Reversible Compartments Enriched in Damaged DNA. Cell Rep, 27, 1809–1821 e1805.

36. Machour, F.E., Barisaac, A.S. and Ayoub, N. (2025) Why are RNA processing factors recruited to DNA double-strand breaks? Trends Genet, 41, 194–200.

37. Marinello, J., Chillemi, G., Bueno, S., Manzo, S.G. and Capranico, G. (2013) Antisense transcripts enhanced by camptothecin at divergent CpG-island promoters associated with bursts of topoisomerase I-DNA cleavage complex and R-loop formation. Nucleic Acids Res, 41, 10110–10123.

38. Valenzisi, P., Marabitti, V., Pichierri, P. and Franchitto, A. (2024) WRNIP1 prevents transcription-associated genomic instability. Elife, 12.

39. Crossley, M.P., Brickner, J.R., Song, C., Zar, S.M.T., Maw, S.S., Chedin, F., Tsai, M.S. and Cimprich, K.A. (2021) Catalytically inactive, purified RNase H1: A specific and sensitive probe for RNA-DNA hybrid imaging. J Cell Biol, 220.

40. Cristini, A., Groh, M., Kristiansen, M.S. and Gromak, N. (2018) RNA/DNA Hybrid Interactome Identifies DXH9 as a Molecular Player in Transcriptional Termination and R-Loop-Associated DNA Damage. Cell Rep, 23, 1891–1905.

41. Wang, K., Wang, H., Li, C., Yin, Z., Xiao, R., Li, Q., Xiang, Y., Wang, W., Huang, J., Chen, L. et al. (2021) Genomic profiling of native R loops with a DNA-RNA hybrid recognition sensor. Sci Adv, 7.

42. Semlow, D.R., Blanco, M.R., Walter, N.G. and Staley, J.P. (2016) Spliceosomal DEAH-Box ATPases Remodel Pre-mRNA to Activate Alternative Splice Sites. Cell, 164, 985–998.

43. Xie, Q., Liu, J., Shan, Y., Wang, S. and Liu, F. (2018) Substrate Determinants for Unwinding Activity of the DExH/D-Box Protein RNA Helicase A. Biochemistry, 57, 6662–6668.

44. Lim, G., Hwang, S., Yu, K., Kang, J.Y., Kang, C. and Hohng, S. (2023) Translocating RNA polymerase generates R-loops at DNA double-strand breaks without any additional factors. Nucleic Acids Res, 51, 9838–9848.

45. Kruger, A., Burkle, A., Hauser, K. and Mangerich, A. (2020) Real-time monitoring of PARP1-dependent PARylation by ATR-FTIR spectroscopy. Nat Commun, 11, 2174.

46. Garcia-Muse, T. and Aguilera, A. (2016) Transcription-replication conflicts: how they occur and how they are resolved. Nat Rev Mol Cell Biol, 17, 553–563.

47. Lin, R., Zhong, X., Zhou, Y., Geng, H., Hu, Q., Huang, Z., Hu, J., Fu, X.D., Chen, L. and Chen, J.Y. (2022) R-loopBase: a knowledgebase for genome-wide R-loop formation and regulation. Nucleic Acids Res, 50, D303–D315.

48. Calame, D.G., Guo, T., Wang, C., Garrett, L., Jolly, A., Dawood, M., Kurolap, A., Henig, N.Z., Fatih, J.M., Herman, I. et al. (2023) Monoallelic variation in DHX9, the gene encoding the DExH-box helicase DHX9, underlies neurodevelopment disorders and Charcot-Marie-Tooth disease. Am J Hum Genet, 110, 1394–1413.

49. Rizzotto, D., Zaccara, S., Rossi, A., Galbraith, M.D., Andrysik, Z., Pandey, A., Sullivan, K.D., Quattrone, A., Espinosa, J.M., Dassi, E. et al. (2020) Nutlin-Induced Apoptosis Is Specified by a Translation Program Regulated by PCBP2 and DHX30. Cell Rep, 30, 4355–4369 e4356.

50. Hikiami, R., Morimura, T., Ayaki, T., Tsukiyama, T., Morimura, N., Kusui, M., Wada, H., Minamiyama, S., Shodai, A., Asada-Utsugi, M. et al. (2022) Conformational change of RNA-helicase DHX30 by ALS/FTD-linked FUS induces mitochondrial dysfunction and cytosolic aggregates. Sci Rep, 12, 16030.

51. Crossley, M.P., Bocek, M. and Cimprich, K.A. (2019) R-Loops as Cellular Regulators and Genomic Threats. Mol Cell, 73, 398–411.

52. Turinetto, V. and Giachino, C. (2015) Multiple facets of histone variant H2AX: a DNA double-strand-break marker with several biological functions. Nucleic Acids Res, 43, 2489–2498.

53. Shibata, A. and Jeggo, P.A. (2020) Roles for 53BP1 in the repair of radiation-induced DNA double strand breaks. DNA Repair (Amst), 93, 102915.

54. Wu, X. and Wang, B. (2021) Abraxas suppresses DNA end resection and limits break-induced replication by controlling SLX4/MUS81 chromatin loading in response to TOP1 inhibitor-induced DNA damage. Nat Commun, 12, 4373.

55. Olive, P.L. and Banath, J.P. (2006) The comet assay: a method to measure DNA damage in individual cells. Nat Protoc, 1, 23–29.

56. Heddle, J.A. (1972) A rapid in vivo test for chromosomal damage. Mutation Research, 18.

57. Her, J., Ray, C., Altshuler, J., Zheng, H. and Bunting, S.F. (2018) 53BP1 Mediates ATR-Chk1 Signaling and Protects Replication Forks under Conditions of Replication Stress. Mol Cell Biol, 38.

58. Amon, J.D. and Koshland, D. (2016) RNase H enables efficient repair of R-loop induced DNA damage. Elife, 5.

59. Bahrami, S. and Drablos, F. (2016) Gene regulation in the immediate-early response process. Adv Biol Regul, 62, 37–49.

60. Tullai, J.W., Schaffer, M.E., Mullenbrock, S., Sholder, G., Kasif, S. and Cooper, G.M. (2007) Immediate-early and delayed primary response genes are distinct in function and genomic architecture. J Biol Chem, 282, 23981–23995.

61. Kumar, C. and Remus, D. (2024) Looping out of control: R-loops in transcription-replication conflict. Chromosoma, 133, 37–56.

62. Guo, H., Golczer, G., Wittner, B.S., Langenbucher, A., Zachariah, M., Dubash, T.D., Hong, X., Comaills, V., Burr, R., Ebright, R.Y. et al. (2021) NR4A1 regulates expression of immediate early genes, suppressing replication stress in cancer. Mol Cell, 81, 4041–4058 e4015.

63. Becker, R.A. and Hub, J.S. (2023) Molecular simulations of DEAH-box helicases reveal control of domain flexibility by ligands: RNA, ATP, ADP, and G-patch proteins. Biol Chem, 404, 867–879.

64. Tauchert, M.J., Fourmann, J.B., Luhrmann, R. and Ficner, R. (2017) Structural insights into the mechanism of the DEAH-box RNA helicase Prp43. Elife, 6.

65. Hamann, F., Enders, M. and Ficner, R. (2019) Structural basis for RNA translocation by DEAH-box ATPases. Nucleic Acids Res, 47, 4349–4362.

66. Zardoni, L., Nardini, E., Brambati, A., Lucca, C., Choudhary, R., Loperfido, F., Sabbioneda, S. and Liberi, G. (2021) Elongating RNA polymerase II and RNA:DNA hybrids hinder fork progression and gene expression at sites of head-on replication-transcription collisions. Nucleic Acids Res, 49, 12769–12784.

67. Zatreanu, D., Han, Z., Mitter, R., Tumini, E., Williams, H., Gregersen, L., Dirac-Svejstrup, A.B., Roma, S., Stewart, A., Aguilera, A. et al. (2019) Elongation Factor TFIIS Prevents Transcription Stress and R-Loop Accumulation to Maintain Genome Stability. Mol Cell, 76, 57–69 e59.

68. Nudler, E. (2012) RNA polymerase backtracking in gene regulation and genome instability. Cell, 149, 1438–1445.

69. Cargill, M., Venkataraman, R. and Lee, S. (2021) DEAD-Box RNA Helicases and Genome Stability. Genes (Basel), 12.

70. Miyamoto, Y., Saiwaki, T., Yamashita, J., Yasuda, Y., Kotera, I., Shibata, S., Shigeta, M., Hiraoka, Y., Haraguchi, T. and Yoneda, Y. (2004) Cellular stresses induce the nuclear accumulation of importin alpha and cause a conventional nuclear import block. J Cell Biol, 165, 617–623.

71. Kose, S. and Imamoto, N. (2014) Nucleocytoplasmic transport under stress conditions and its role in HSP70 chaperone systems. Biochim Biophys Acta, 1840, 2953–2960.

72. Lin, W.L., Chen, J.K., Wen, X., He, W., Zarceno, G.A., Chen, Y., Chen, S., Paull, T.T. and Liu, H.W. (2022) DDX18 prevents R-loop-induced DNA damage and genome instability via PARP-1. Cell Rep, 40, 111089.

73. Boleslavska, B., Oravetzova, A., Shukla, K., Nascakova, Z., Ibini, O.N., Hasanova, Z., Andrs, M., Kanagaraj, R., Dobrovolna, J. and Janscak, P. (2022) DDX17 helicase promotes resolution of R-loop-mediated transcription-replication conflicts in human cells. Nucleic Acids Res, 50, 12274–12290.

74. Song, C., Hotz-Wagenblatt, A., Voit, R. and Grummt, I. (2017) SIRT7 and the DEAD-box helicase DDX21 cooperate to resolve genomic R loops and safeguard genome stability. Genes Dev, 31, 1370–1381.

75. Chakraborty, P., Huang, J.T.J. and Hiom, K. (2018) DHX9 helicase promotes R-loop formation in cells with impaired RNA splicing. Nat Commun, 9, 4346.

76. Secchi, M., Garbelli, A., Riva, V., Deidda, G., Santonicola, C., Formica, T.M., Sabbioneda, S., Crespan, E. and Maga, G. (2024) Synergistic action of human RNaseH2 and the RNA helicase-nuclease DDX3X in processing R-loops. Nucleic Acids Res, 52, 11641–11658.

77. Wang, I.X., Grunseich, C., Fox, J., Burdick, J., Zhu, Z., Ravazian, N., Hafner, M. and Cheung, V.G. (2018) Human proteins that interact with RNA/DNA hybrids. Genome Res, 28, 1405–1414.

78. Hamperl, S., Bocek, M.J., Saldivar, J.C., Swigut, T. and Cimprich, K.A. (2017) Transcription-Replication Conflict Orientation Modulates R-Loop Levels and Activates Distinct DNA Damage Responses. Cell, 170, 774–786 e719.

79. Petermann, E., Lan, L. and Zou, L. (2022) Sources, resolution and physiological relevance of R-loops and RNA-DNA hybrids. Nat Rev Mol Cell Biol, 23, 521–540.

80. Cristini, A., Tellier, M., Constantinescu, F., Accalai, C., Albulescu, L.O., Heiringhoff, R., Bery, N., Sordet, O., Murphy, S. and Gromak, N. (2022) RNase H2, mutated in Aicardi-Goutieres syndrome, resolves co-transcriptional R-loops to prevent DNA breaks and inflammation. Nat Commun, 13, 2961.

81. Khan, Z.U., Carretero-Rey, M., de Leon-Lopez, C.A.M. and Navarro-Lobato, I. (2025) Memory-Associated Immediate Early Genes: Roles in Synaptic Function, Memory Processes, and Neurological Diseases. Mol Neurobiol, 62, 15885–15915.

82. Matos, D.A., Zhang, J.M., Ouyang, J., Nguyen, H.D., Genois, M.M. and Zou, L. (2020) ATR Protects the Genome against R Loops through a MUS81-Triggered Feedback Loop. Mol Cell, 77, 514–527 e514.

83. Murai, J., Zhang, H., Pongor, L., Tang, S.W., Jo, U., Moribe, F., Ma, Y., Tomita, M. and Pommier, Y. (2020) Chromatin Remodeling and Immediate Early Gene Activation by SLFN11 in Response to Replication Stress. Cell Rep, 30, 4137–4151 e4136.

