## Supplementary figures for "DHX30 is a replication stress-responsive R-loop helicase that safe-guards genome stability"

### Supplementary information:

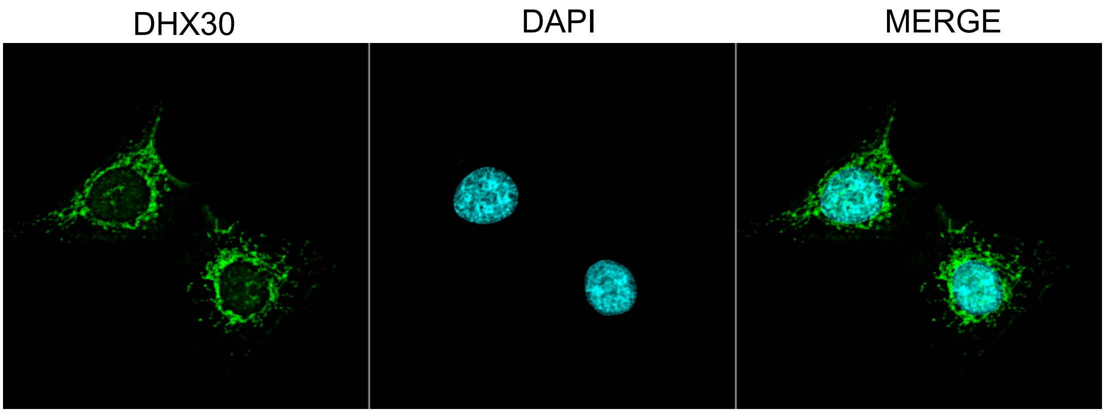


**Supplementary Movie 1. Subcellular localization of endogenous DHX30.** Representative single optical section from the confocal z-stack used to generate Supplementary Movie 1, showing endogenous DHX30 (green), DAPI-stained nuclei (cyan), and the merged image.

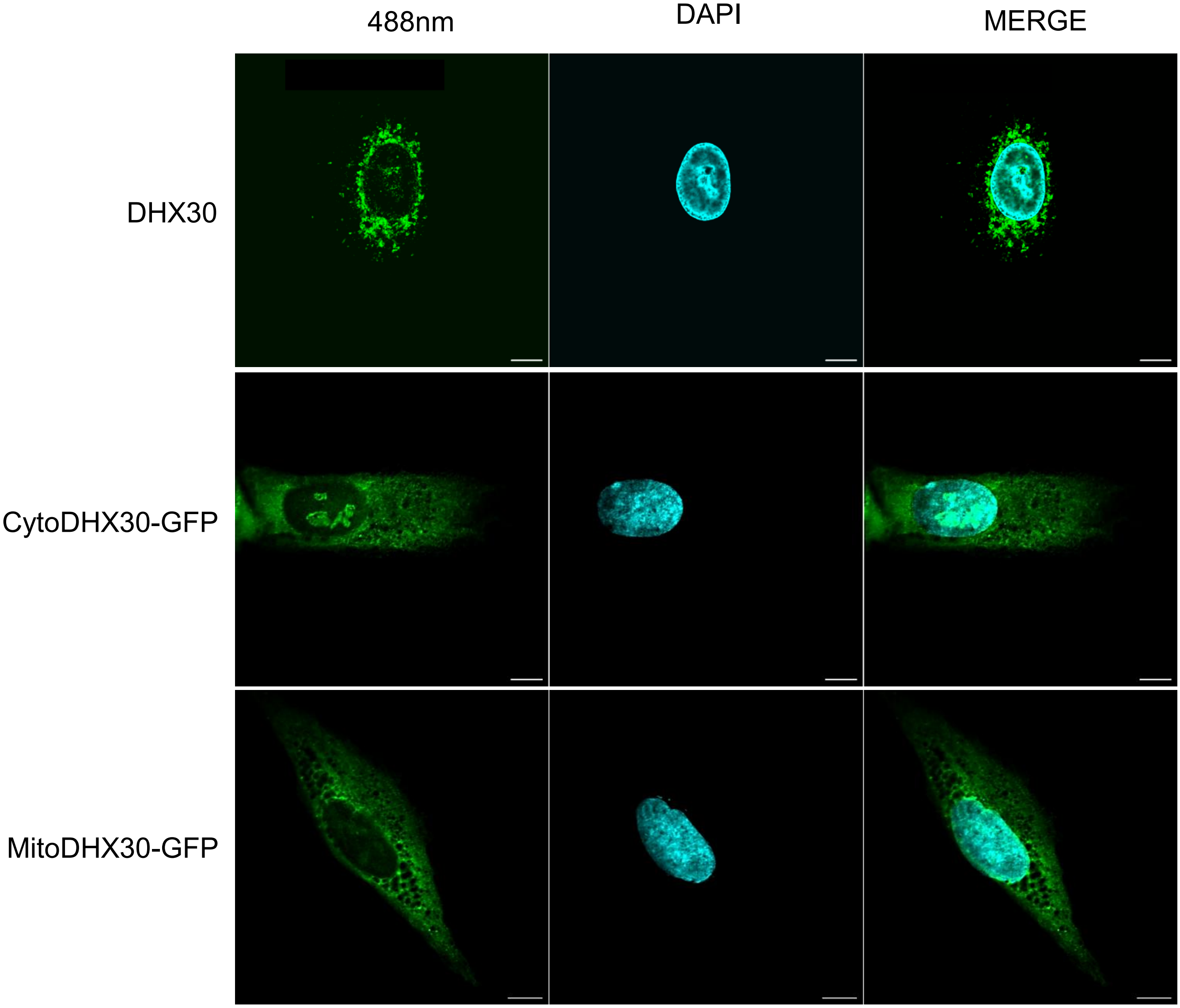


**Supplementary Fig. S1. Subcellular localization of DHX30.** Representative single-plane confocal images showing endogenous DHX30 (top), CytoDHX30-GFP (middle), and MitoDHX30-GFP (bottom). DHX30 Ab/GFP fluorescence is shown in green and nuclei were counterstained with DAPI (cyan). Merged images are shown in the right panels. Scale bars, 10µm.

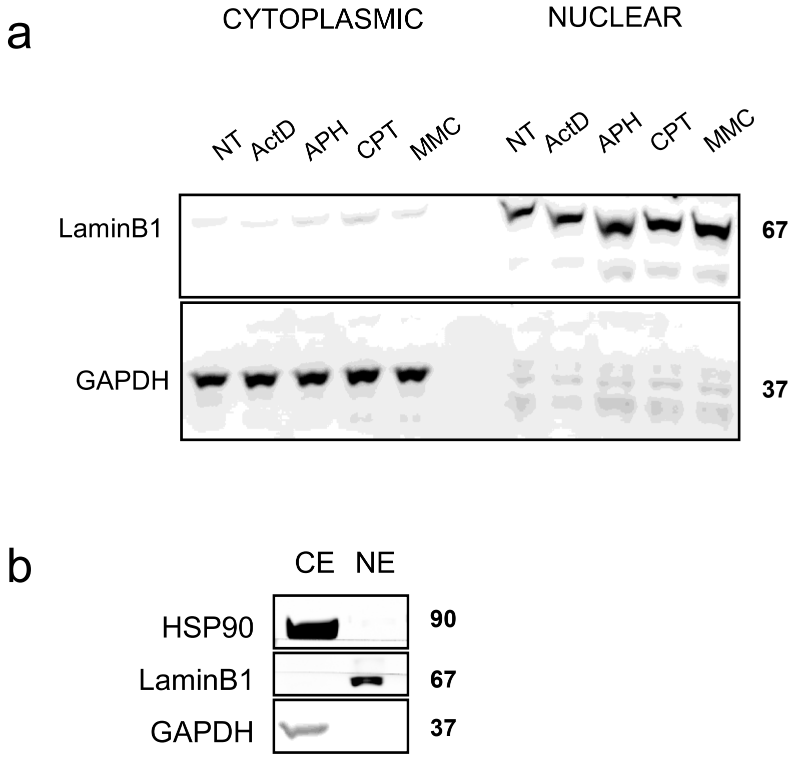


**Supplementary Fig. S2. Validation of cytoplasmic and nuclear fractionation. a** Immunoblot analysis of cytoplasmic and nuclear fractions from HEK293T cells under untreated (NT) conditions or following treatment with actinomycin D (ActD), aphidicolin (APH), camptothecin (CPT), or mitomycin C (MMC). LaminB1 and GAPDH were used as nuclear and cytoplasmic markers, respectively, to assess fractionation purity. Cytoplasmic and nuclear fractions were resolved on the same gel and transferred to the same membrane; the two fractions are shown at the same exposure. **b** Representative immunoblot of cytoplasmic extract (CE) and nuclear extract (NE) showing HSP90 and GAPDH as cytoplasmic markers and Lamin B1 as a nuclear marker. Molecular weights are indicated in kDa.

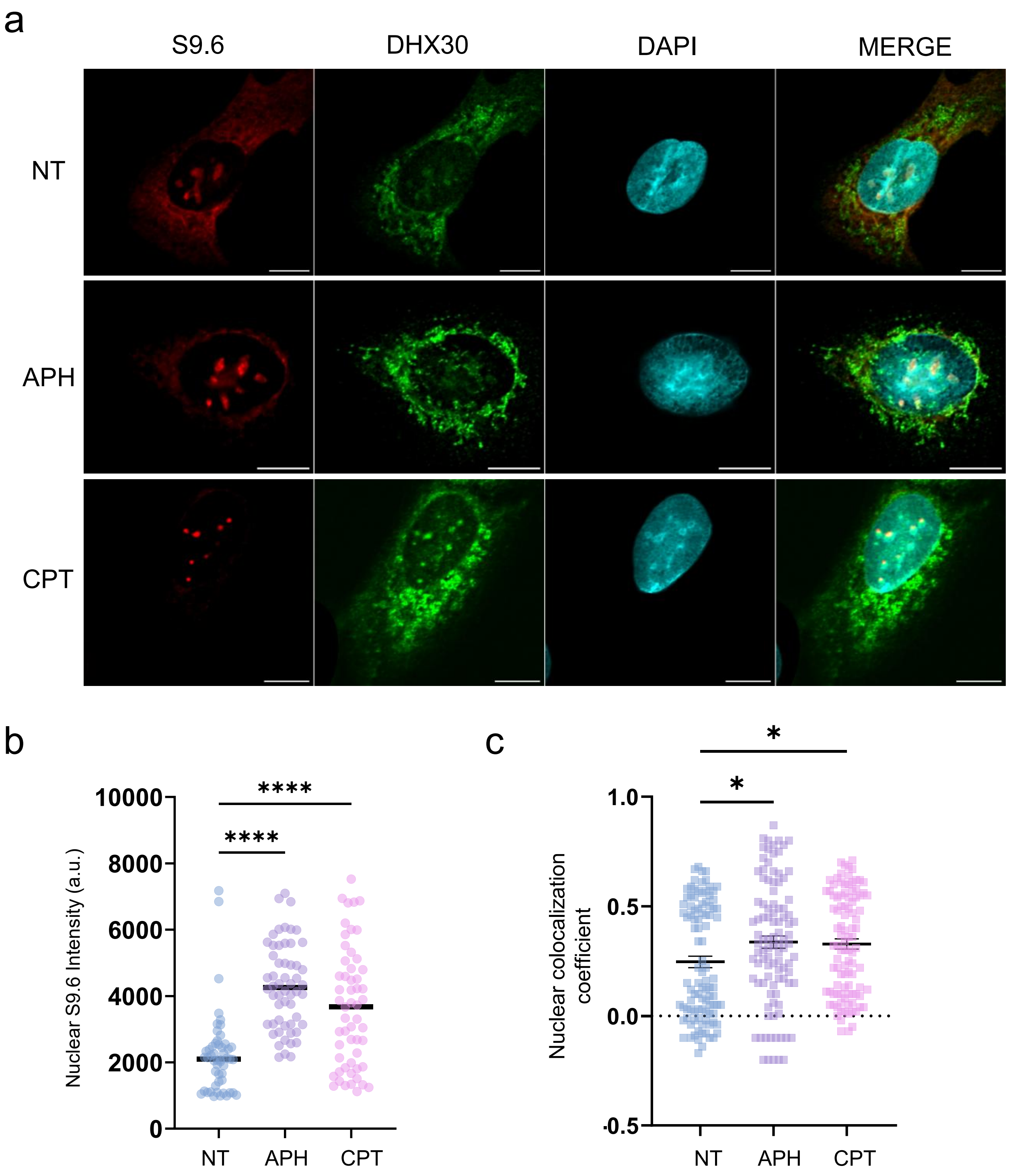


**Supplementary Fig. S3. DHX30 overlap with S9.6-positive nuclear signal increases following APH and CPT treatment. a** Representative single-plane confocal image of untreated (NT) and treated (APH, CPT) U2OS cells stained for endogenous DHX30 (green), S9.6 (red) and DAPI (cyan). Scale bar, 10 µm. **b** Quantification of nuclear S9.6 fluorescence intensity in NT and treated cells (APH,CPT). Each point represents one cell. A total of ~50 cells per condition, pooled from two independent biological experiments, were analyzed. Statistical significance was assessed using ordinary one-way ANOVA followed by Dunnett's multiple-comparisons test, comparing APH- and CPT-treated cells with the NT condition (****P<0,0001). **c** Quantification of nuclear overlap between DHX30 and S9.6-positive signal under NT, APH, and CPT conditions. Pearson's R (no threshold) was measured using the Coloc2 plugin in Fiji. Each point represents one cell. Approximately 100 cells per condition, pooled from three independent biological experiments, were analyzed. Statistical significance was assessed using ordinary one-way ANOVA followed by Dunnett's multiple-comparisons test, comparing APH- and CPT-treated cells with the NT condition (*P < 0.05).

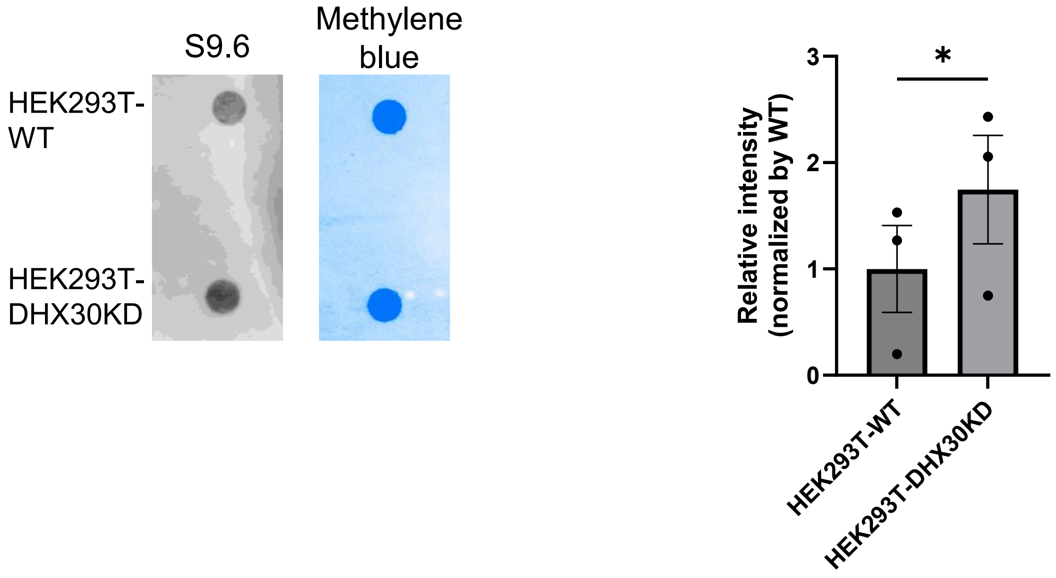


**Supplementary Fig. S4. Normalization of S9.6 signal to the total DNA.** Representative dot blot analysis of S9.6 signal in total DNA isolated from HEK293T-WT and HEK293T-DHX30KD cells, with methylene blue staining used as a loading control for the total DNA. Right, quantification of S9.6 signal intensity normalized to the corresponding methylene blue signal and expressed relative to HEK293T-WT. Data are presented as mean ± SEM from three independent biological experiments. Statistical significance was assessed using a paired two-tailed t-test ($P < 0.05$).

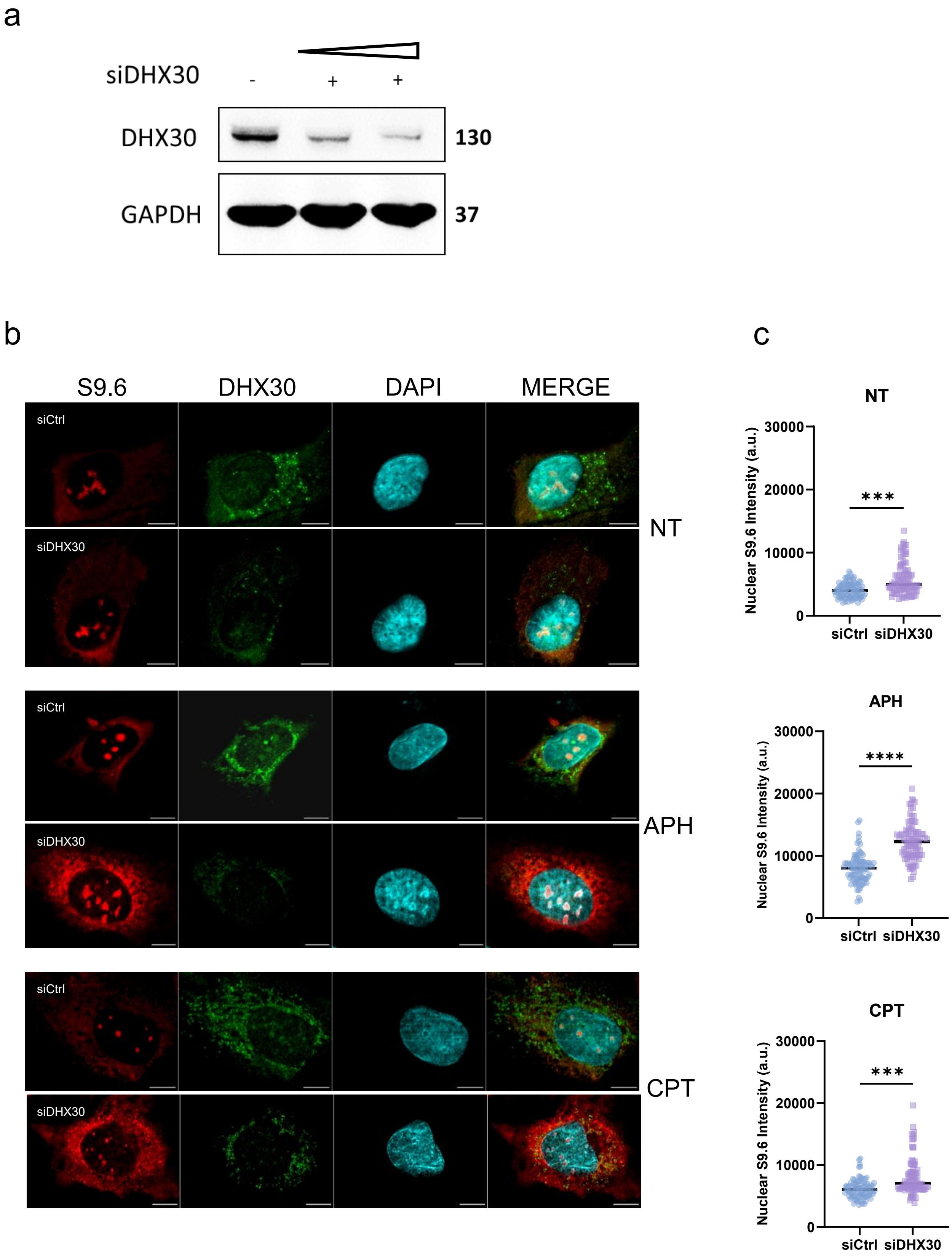

a
siDHX30
-
+
+
DHX30
130
GAPDH
37
b
S9.6
DHX30
DAPI
MERGE
siCtrl
siDHX30
NT
siCtrl
siDHX30
APH
siCtrl
siDHX30
CPT
c
NT
Nuclear S9.6 Intensity (a.u.)
30000
20000
10000
0
***
siCtrl
siDHX30
APH
Nuclear S9.6 Intensity (a.u.)
30000
20000
10000
0
****
siCtrl
siDHX30
CPT
Nuclear S9.6 Intensity (a.u.)
30000
20000
10000
0
***
siCtrl
siDHX30

**Supplementary Fig. S5. Transient DHX30 depletion increases S9.6-positive nuclear signal under basal and replication stress conditions. a** Immunoblot analysis showing the efficiency of transient DHX30 depletion by siRNA. Cells transfected with control siRNA (siCtrl) or increasing amounts of DHX30-targeting siRNA (siDHX30) show reduced DHX30 protein levels. GAPDH was used as a loading control. **b** Representative single-plane confocal images of U2OS cells transfected with siCtrl or siDHX30 under untreated (NT) conditions or following treatment with aphidicolin (APH) or camptothecin (CPT). Cells were stained with S9.6 (red), DHX30 (green), and DAPI (cyan). Scale bar, 10 μm. **c** Quantification of nuclear S9.6 fluorescence intensity in siCtrl and siDHX30 cells under the indicated conditions. Each point represents one cell. Approximately 100 cells per condition, pooled from three independent biological experiments, were analyzed. Statistical significance was assessed using ordinary one-way ANOVA followed by Šídák’s multiple-comparisons test for the indicated pairwise comparisons (**$P < 0.01$; ***$P < 0.001$; ****$P < 0.0001$).

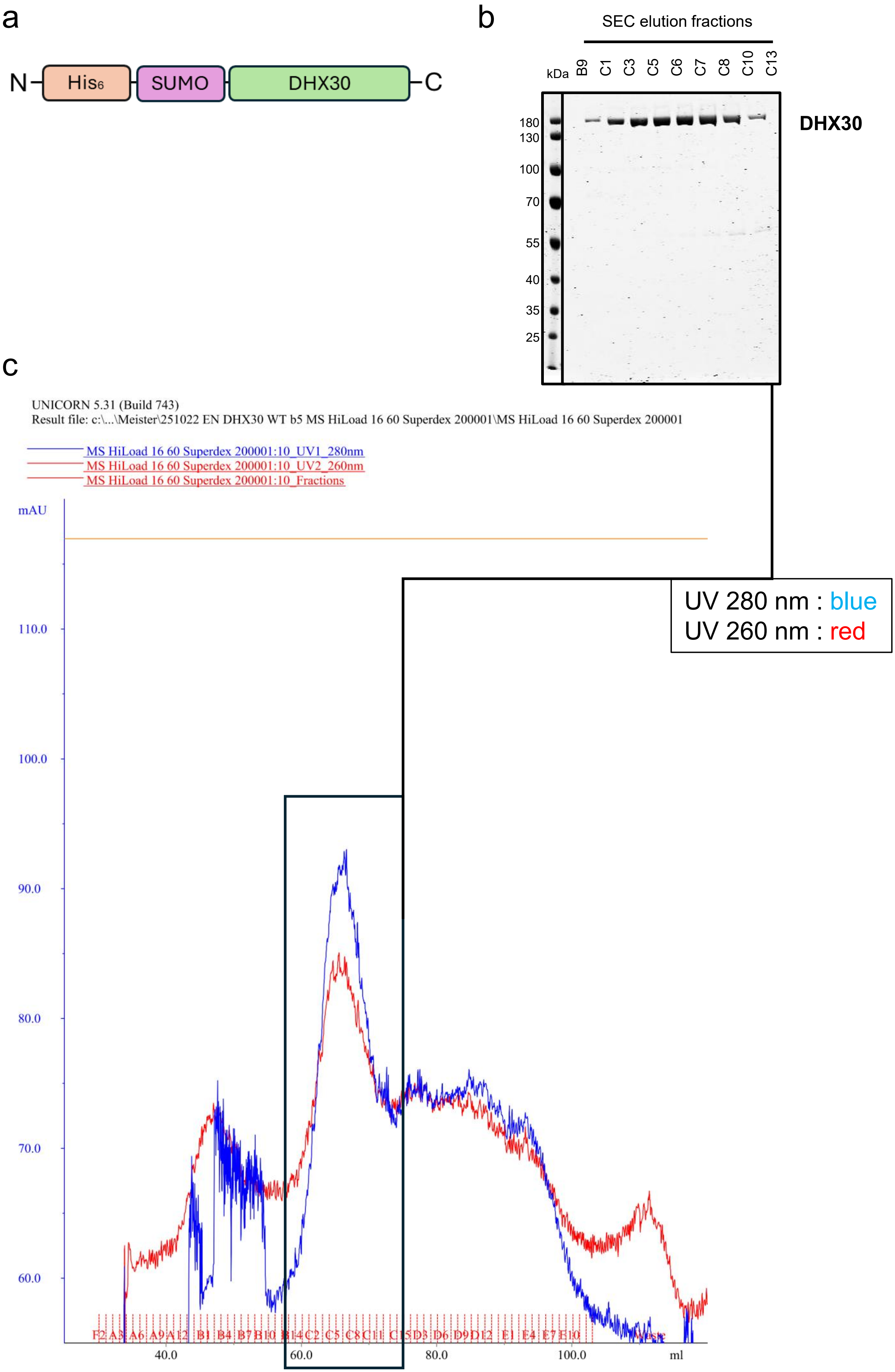

a
N
His6
SUMO
DHX30
C
b
SEC elution fractions
kDa
B9
C1
C3
C5
C6
C7
C8
C10
C13
180
130
100
70
55
40
35
25
DHX30
c
UNICORN 5.31 (Build 743)
Result file: c:\...\Meister\251022 EN DHX30 WT b5 MS HiLoad 16 60 Superdex 200001\MS HiLoad 16 60 Superdex 200001
MS HiLoad 16 60 Superdex 200001:10_UV1_280nm
MS HiLoad 16 60 Superdex 200001:10_UV2_260nm
MS HiLoad 16 60 Superdex 200001:10_Fractions
mAU
110.0
100.0
90.0
80.0
70.0
60.0
40.0
60.0
80.0
100.0
ml
UV 280 nm : blue
UV 260 nm : red

**Supplementary Fig. S6. Purification of recombinant DHX30. a** Schematic representation of the recombinant DHX30 construct carrying an N-terminal $His_6$-SUMO tag. **b** SDS-PAGE analysis of size-exclusion chromatography (SEC) elution fractions of affinity-purified DHX30. Fractions corresponding to the target protein peak are shown. **c** SEC chromatogram of affinity-purified DHX30. The blue and red traces represent UV absorbance at 280 nm and 260 nm, respectively. Fraction positions are indicated along the x-axis. The boxed region indicates the fractions corresponding to the major DHX30-containing peak analyzed in b.

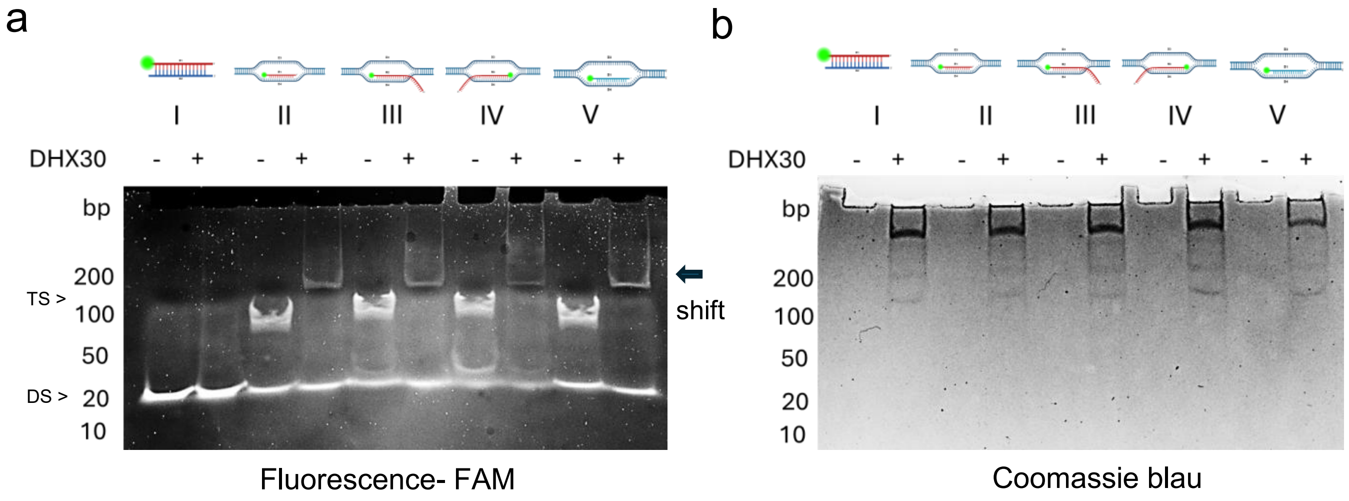


**Supplementary Fig. S7. DHX30 binds three-stranded R-loop and D-loop structures. a** Electrophoretic mobility shift assay (EMSA) using five different FAM-labelled nucleic acid substrates (I–V) in the absence (−) or presence (+) of recombinant DHX30. Formation of slower-migrating species upon addition of DHX30 indicates detectable binding to the indicated substrates. Double-stranded (DS) and three-stranded (TS) species are indicated. **b** Coomassie blue staining of the gel corresponding to the EMSA reactions shown in **a**.

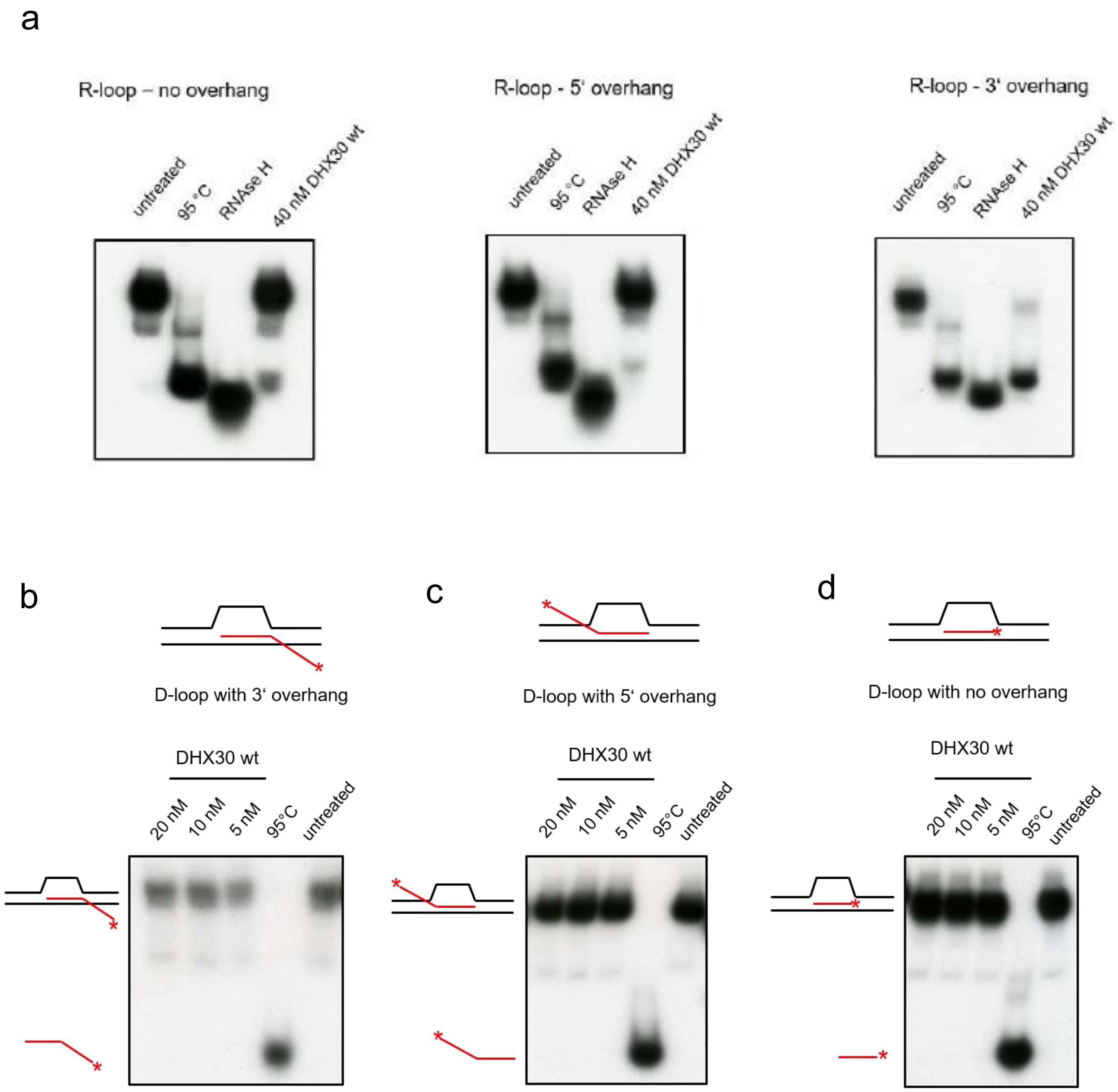


**Supplementary Fig. S8. Validation of R-loop substrates and assessment of DHX30 helicase activity on D-loop substrates. a** RNase H digestion of radiolabelled R-loop substrates (used in Figure 3). Samples lacking overhangs (left), containing a 5′ RNA overhang (middle) and containing a 3′ RNA overhang (right) were incubated under the indicated conditions: untreated samples; heat-denatured samples (95°C); RNase H-treated samples (1 U, 1 h at 37°C); and samples incubated with DHX30 WT (40 nM). The samples were then resolved on a native PAGE. **b-d** Representative helicase assays using wild-type DHX30 and D-loop substrates containing a 3′ overhang (**b**), a 5′ overhang (**c**), or no overhang (**d**). Increasing concentrations of DHX30 (5–20 nM) were incubated with the indicated substrates. Heat-denatured (95 °C) and untreated substrates were included as controls. Reaction products were resolved by native PAGE. Red lines indicate the radiolabelled strand, and red asterisks denote the radiolabelled end.

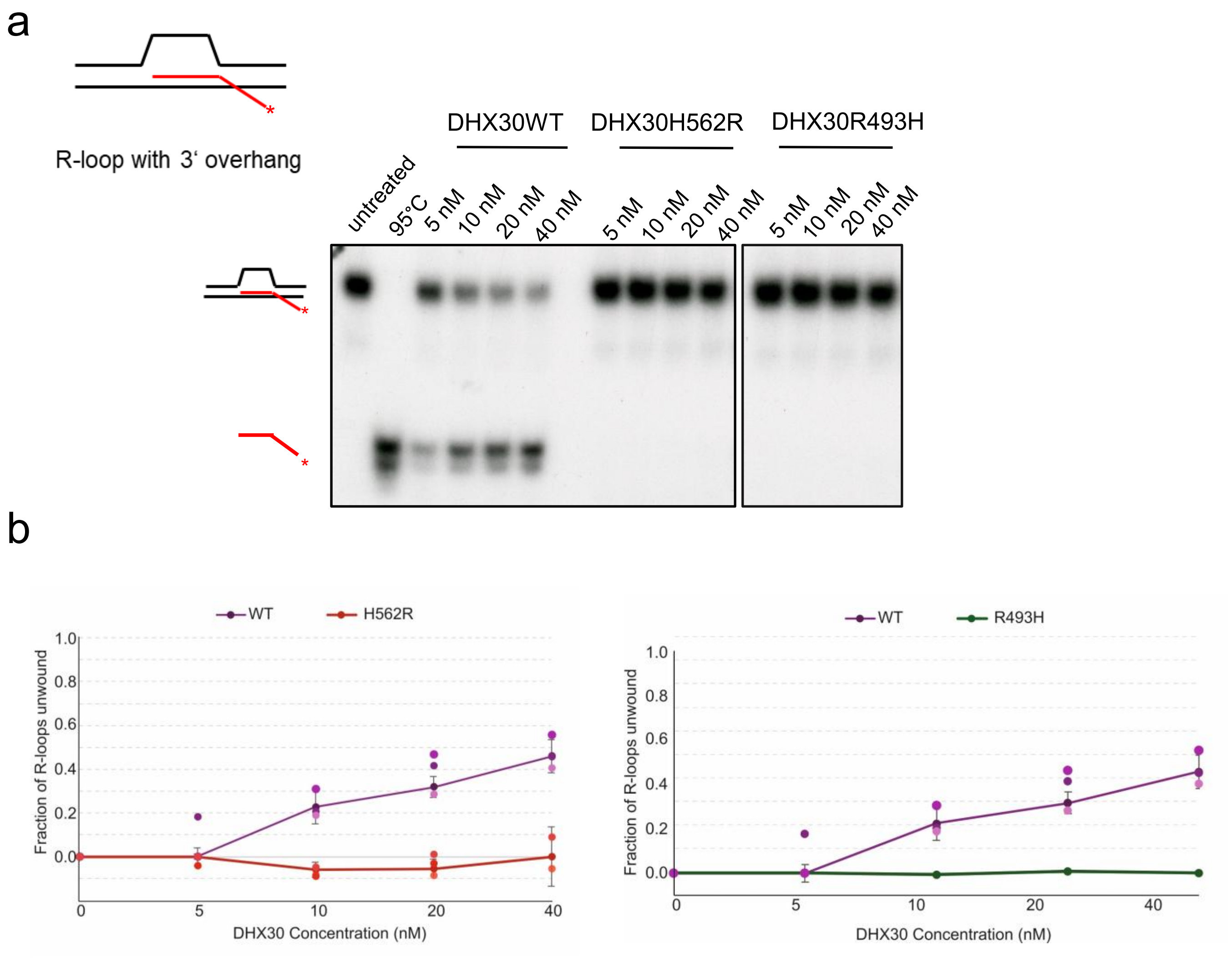


**Supplementary Fig. S9. R-loop unwinding activity of DHX30 WT and disease-associated H562R and R493H variants. a** Representative native PAGE analysis comparing the unwinding of a defined R-loop substrate containing a 3′ single-stranded RNA overhang by wild-type DHX30 (DHX30WT) and the disease-associated DHX30-H562R and DHX30-R493H variants. Reactions were performed with increasing protein concentrations, as indicated. Untreated and heat-denatured (95 °C) substrates were included as controls. The red line indicates the radiolabeled RNA strand, and the asterisk denotes the radiolabeled end. Positions of intact R-loop substrate and released radiolabeled ssRNA are indicated.

**b** Quantification of R-loop unwinding by DHX30WT, DHX30-H562R and DHX30-R493H variants at the indicated protein concentrations. Data are presented as mean ± SEM from three independent experiments. R493H was analyzed in a single independent experiment and is therefore shown descriptively without statistical comparison.

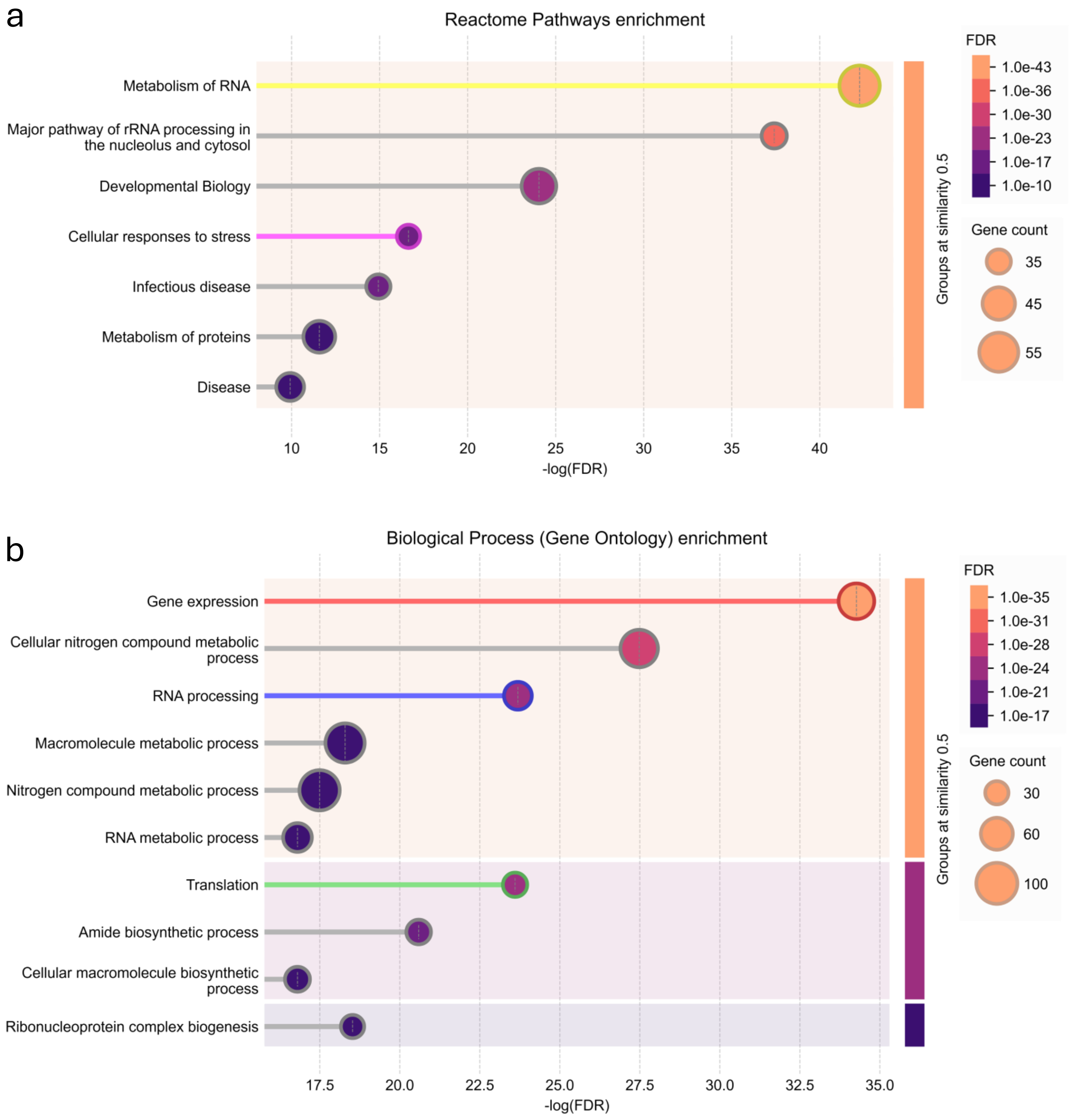


**Supplementary Fig. S10. Mass spectrometry analysis of the DHX30-associated nuclear protein network. a** Reactome pathway and **b** Gene Ontology (GO) biological process enrichment analyses of the DHX30-associated proteins identified by nuclear immunoprecipitation followed by mass spectrometry. Bubble size indicates the number of proteins associated with each term, while colour intensity reflects statistical significance (FDR). Data were generated using the STRING online tool (https://string-db.org/).

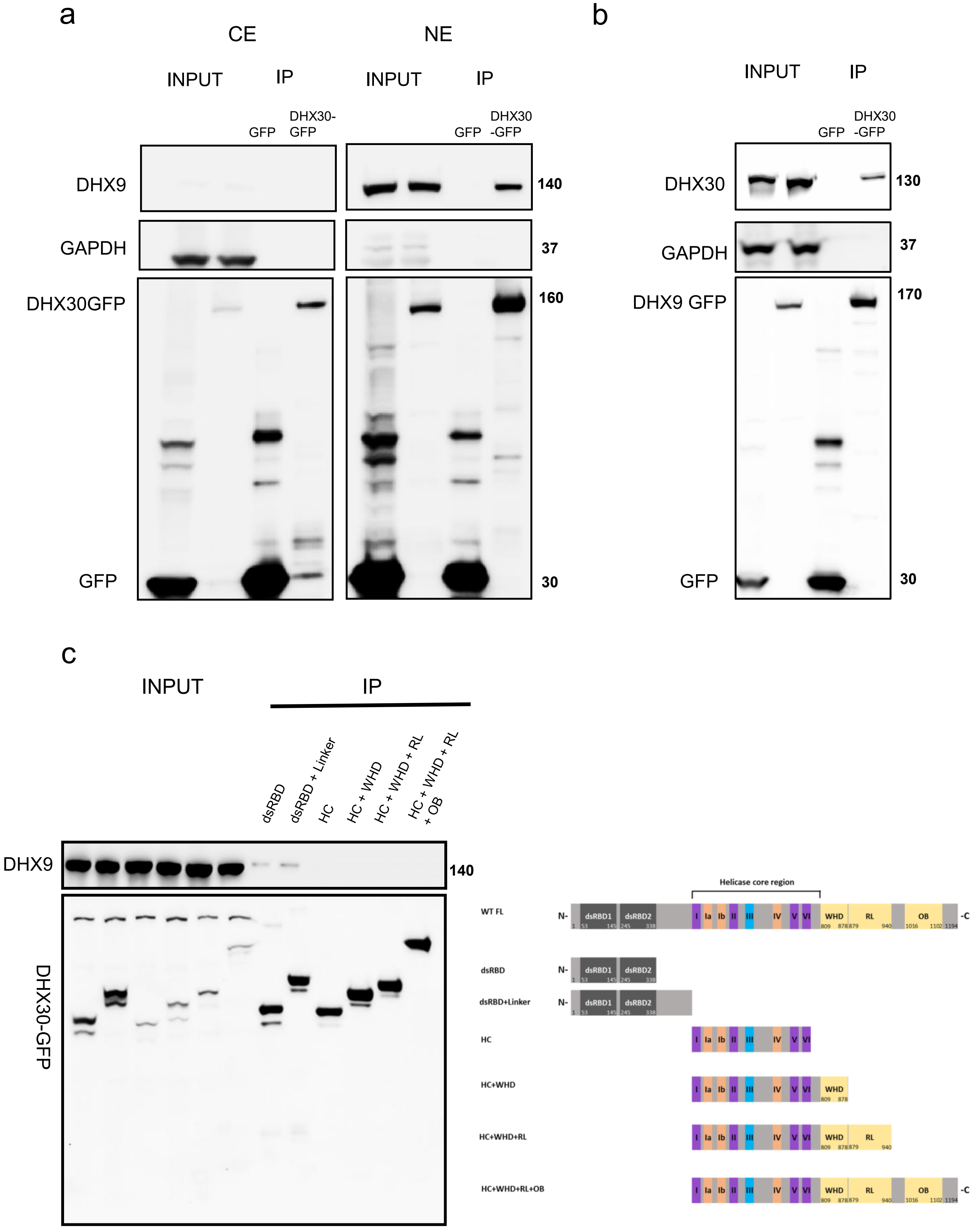
a
CE
NE
INPUT
IP
INPUT
IP
GFP
DHX30-GFP
GFP
DHX30 -GFP
DHX9
140
GAPDH
37
DHX30GFP
160
GFP
30
b
INPUT
IP
GFP
DHX30 -GFP
DHX30
130
GAPDH
37
DHX9 GFP
170
GFP
30
c
INPUT
IP
dsRBD
dsRBD + Linker
HC
HC + WHD
HC + WHD + RL
HC + WHD + RL + OB
DHX9
140
DHX30-GFP
Helicase core region
WT FL
dsRBD
dsRBD+Linker
HC
HC+WHD
HC+WHD+RL
HC+WHD+RL+OB
N-
-C
dsRBD1
dsRBD2
I
Ia
Ib
II
III
IV
V
VI
WHD
RL
OB

**Supplementary Fig. S11. Analysis of the association between DHX30 and DHX9. a** GFP-Trap® immunoprecipitation from cytoplasmic extracts (CE) and nuclear extracts (NE) of cells expressing DHX30-GFP or GFP alone, followed by immunoblot analysis of endogenous DHX9. GAPDH was used as a cytoplasmic marker to assess cellular fractionation. **b** Reciprocal GFP-Trap® immunoprecipitation from whole-cell lysates of cells expressing DHX9-GFP or GFP alone, followed by immunoblot analysis of endogenous DHX30. **c** GFP-Trap® immunoprecipitation of GFP-tagged DHX30 truncation constructs followed by immunoblot analysis of endogenous DHX9 to assess DHX9 co-purification with the different DHX30 constructs. Right, schematic representation of full-length DHX30 and the truncation constructs used in the assay.

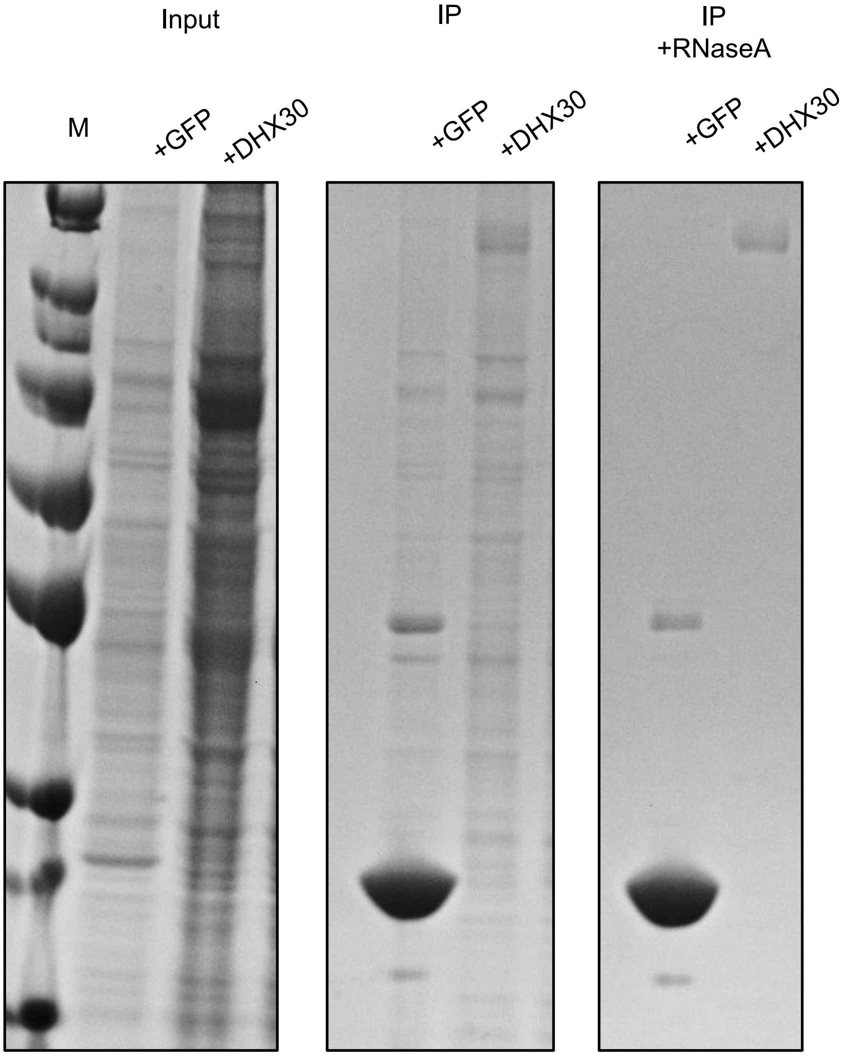


**Supplementary Fig. S12. RNase A treatment reduces proteins co-purifying with DHX30-GFP.** Representative Coomassie-stained SDS-PAGE analysis of GFP-Trap immunoprecipitations from whole-cell lysates expressing GFP alone or DHX30-GFP. Input samples and immunoprecipitated fractions (IP) are shown. Where indicated, samples were treated with RNase A (150 mM NaCl) prior to immunoprecipitation. RNase A treatment markedly reduced the overall pattern of proteins co-purifying with DHX30-GFP, consistent with a substantial RNA-dependent component of DHX30-associated complexes. M, molecular-weight marker.

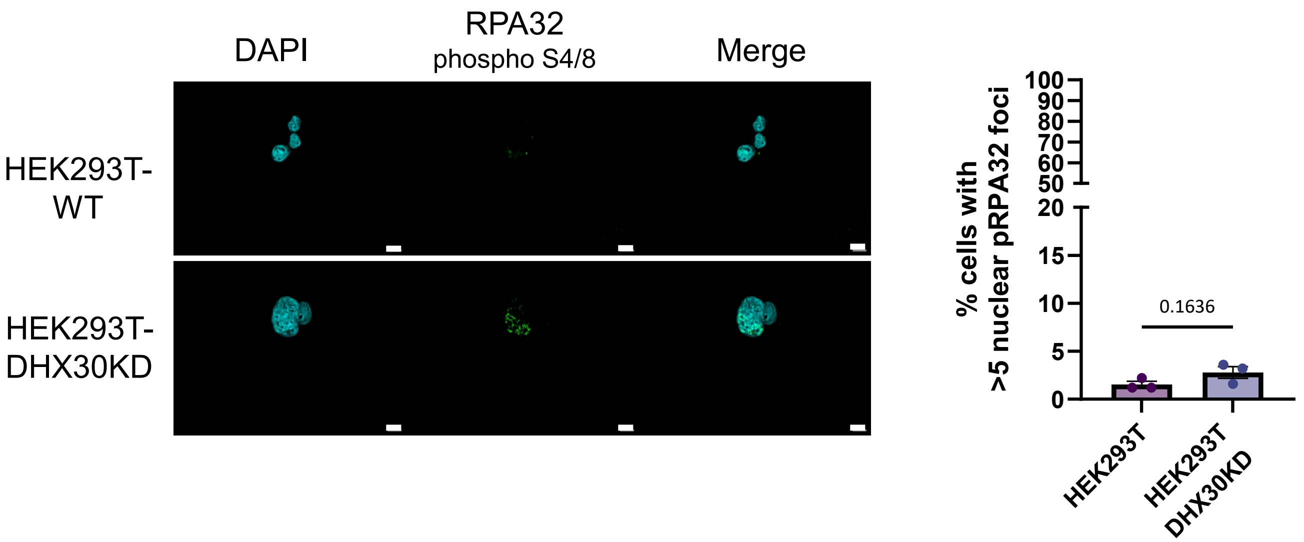


**Supplementary Fig. S13. Analysis of RPA32 S4/S8 phosphorylation in DHX30-deficient cells.** Representative single-plane immunofluorescence images of HEK293T-WT and HEK293T-DHX30KD cells stained for phosphorylated RPA32 S4/S8 (green) and DAPI (cyan). Merged images are shown in the right panels. Right, quantification of the percentage of cells containing >5 nuclear pRPA32 S4/S8 foci. Data are presented as mean ± SEM from three independent biological experiments. Statistical significance was assessed by two-tailed t-test and the difference was not statistically significant (ns) ($P = 0.1636$). Scale bar, 10 μm.

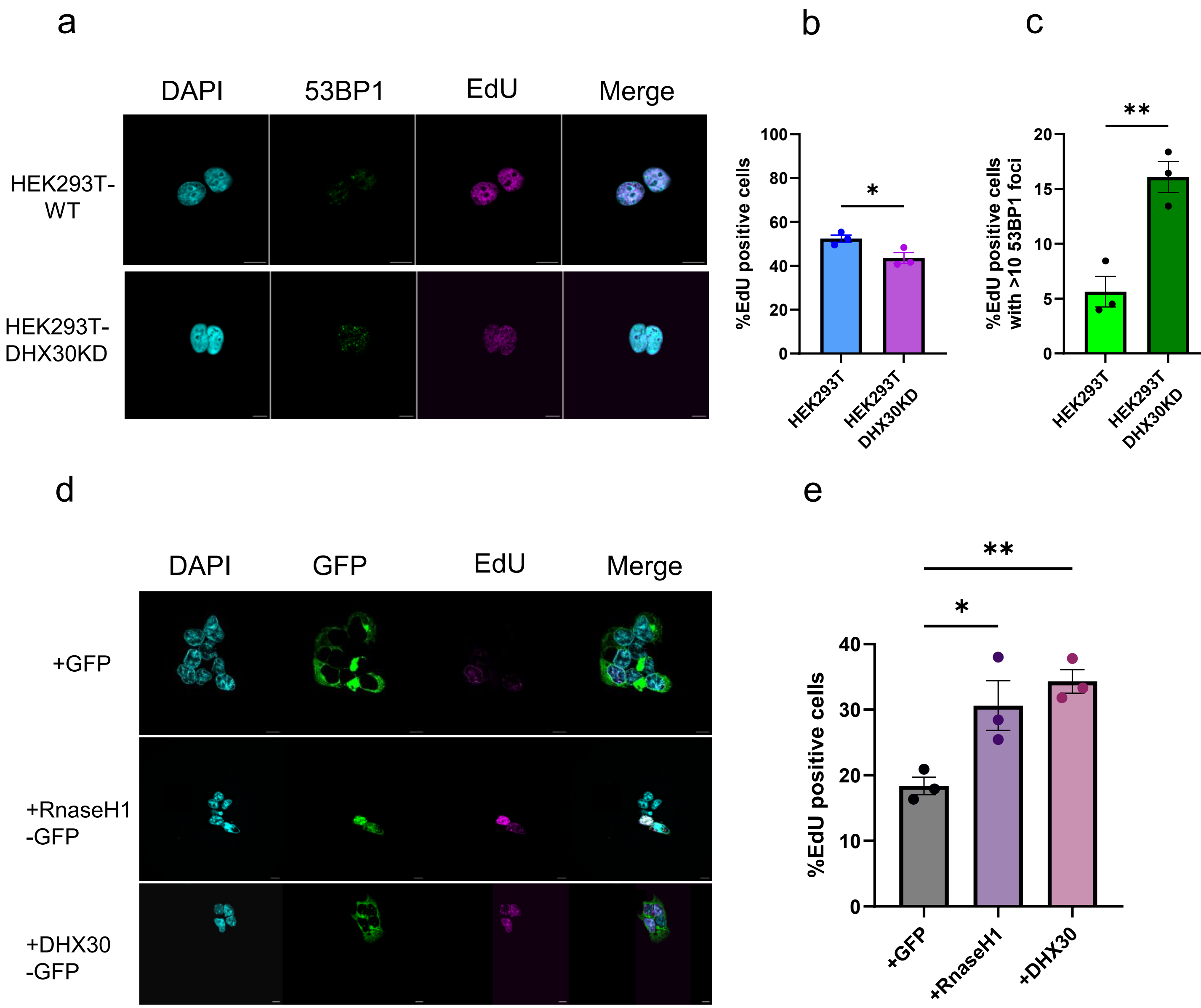
a
DAPI
53BP1
EdU
Merge
HEK293T-WT
HEK293T-DHX30KD
b
%EdU positive cells
100
80
60
40
20
0
*
HEK293T
HEK293T DHX30KD
c
%EdU positive cells with >10 53BP1 foci
20
15
10
5
0
**
HEK293T
HEK293T DHX30KD
d
DAPI
GFP
EdU
Merge
+GFP
+RnaseH1 -GFP
+DHX30 -GFP
e
%EdU positive cells
40
30
20
10
0
**
*
+GFP
+RnaseH1
+DHX30

**Supplementary Fig. S14. DHX30 deficiency reduces EdU incorporation and this phenotype is rescued by RNase H1 and wild-type DHX30 overexpression. a** Representative single-plane immunofluorescence images of HEK293T-WT and HEK293T-DHX30KD cells stained for 53BP1 (green), EdU (magenta), and DAPI (cyan). Merged images are shown in the right panels. **b** Quantification of the percentage of EdU-positive cells in HEK293T-WT and HEK293T-DHX30KD cells. DHX30-deficient cells showed a reduced fraction of EdU-positive cells compared with WT cells. Data are presented as mean ± SEM from three independent biological experiments. **c** Quantification of the percentage of EdU-positive cells containing ≥10 nuclear 53BP1 foci in HEK293T-WT and HEK293T-DHX30KD cells. Data are presented as mean ± SEM from three independent biological experiments. Statistical significance was assessed using an unpaired two-tailed t-test (*$P < 0.05$; **$P < 0.01$;). **d** Representative single-plane immunofluorescence images of DHX30-deficient cells expressing GFP alone, RNase H1-GFP, or wild-type DHX30-GFP. GFP fluorescence is shown in green, EdU incorporation in magenta, and DAPI in cyan. **e** Quantification of the percentage of EdU-positive cells under the conditions shown in **d**. Data are presented as mean ± SEM from three independent biological experiments. Statistical significance was assessed using ordinary one-way ANOVA followed by Dunnett's multiple-comparisons test, comparing each condition with the GFP control (*$P < 0.05$; **$P < 0.01$). Scale bar, 10 μm.

a

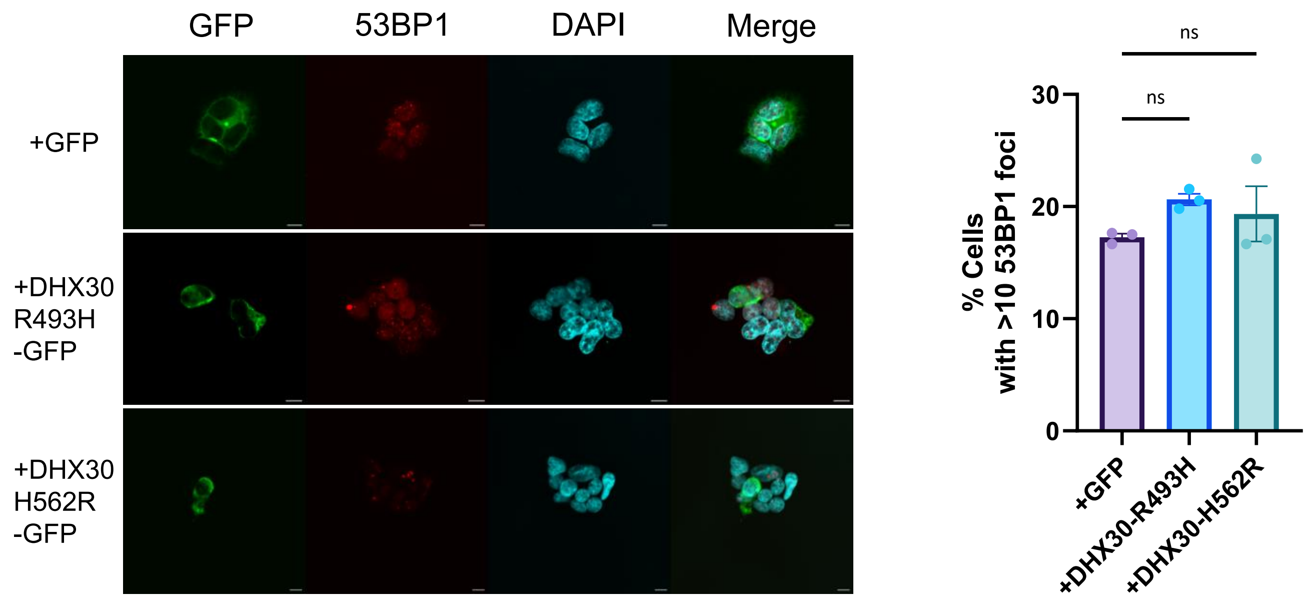
GFP
53BP1
DAPI
Merge
+GFP
+DHX30 R493H -GFP
+DHX30 H562R -GFP
ns
ns
30
20
10
0
% Cells with >10 53BP1 foci
+GFP
+DHX30-R493H
+DHX30-H562R

b

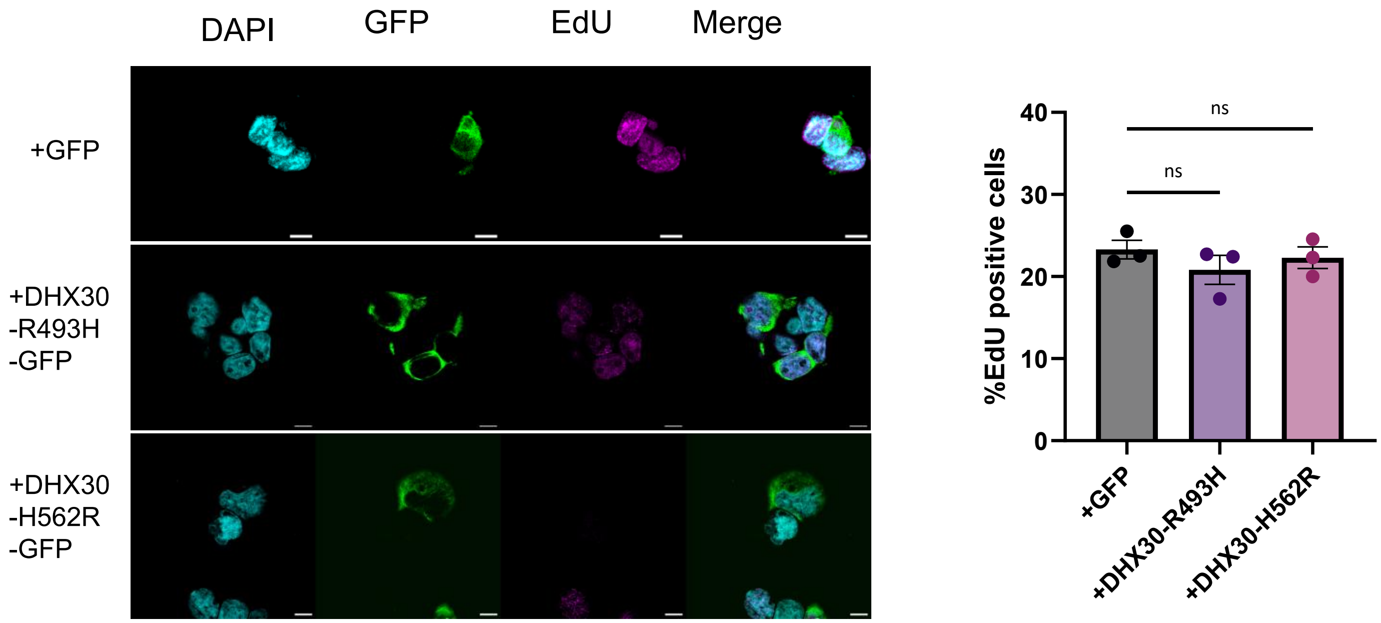
DAPI
GFP
EdU
Merge
+GFP
+DHX30 -R493H -GFP
+DHX30 -H562R -GFP
ns
ns
40
30
20
10
0
%EdU positive cells
+GFP
+DHX30-R493H
+DHX30-H562R

c

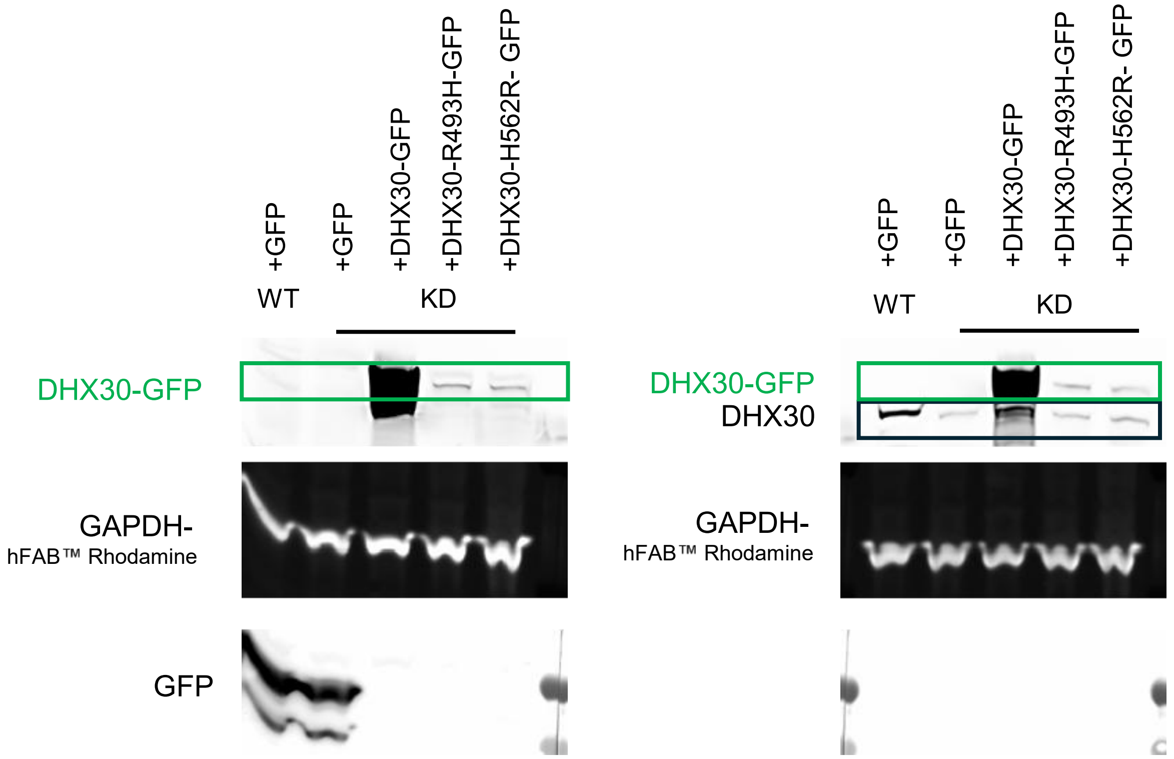


**Supplementary Fig. S15. Disease-associated DHX30 variants fail to significantly reduce 53BP1 foci in DHX30-deficient cells. a** Representative single-plane immunofluorescence images of DHX30-deficient cells expressing GFP alone, DHX30-R493H-GFP, or DHX30-H562R-GFP. Cells were stained for 53BP1 (red) and DAPI (cyan), while GFP fluorescence is shown in green. Merged images are shown in the right panels. Scale bars 10µm. Right, quantification of the percentage of cells containing >10 nuclear 53BP1 foci under the indicated conditions. Data are presented as mean ± SEM from three independent biological experiments. Statistical significance was assessed using ordinary one-way ANOVA followed by Dunnett's multiple-comparisons test, comparing each DHX30 variant with the GFP control (ns, not significant). **b** Representative single-plane immunofluorescence images of DHX30-deficient cells expressing GFP alone, DHX30-R493H-GFP, or DHX30-H562R-GFP. Quantification of the percentage of EdU-positive cells. Statistical significance was assessed using ordinary one-way ANOVA followed by Dunnett's multiple-comparisons test, comparing each DHX30 variant with the GFP control (ns, not significant). **c** Expression of ectopic DHX30 constructs in DHX30-deficient cells. Representative Western blot of HEK293T-WT (WT) and HEK293T-DHX30KD (KD) cells expressing GFP only and HEK293T-DHX30KD (KD) cells expressing wild-type DHX30-GFP (DHX30-WT), or the disease-associated variants DHX30-R493H-GFP and DHX30-H562R-GFP. The membrane was probed with an anti-GFP antibody to verify ectopic expression of the GFP-tagged DHX30 constructs. On the right, the same membrane subsequently probed with an anti-DHX30 antibody is shown to assess endogenous DHX30 levels and confirm DHX30 depletion in DHX30KD cells. GAPDH was used as a loading control. Scale bar, 10 µm.

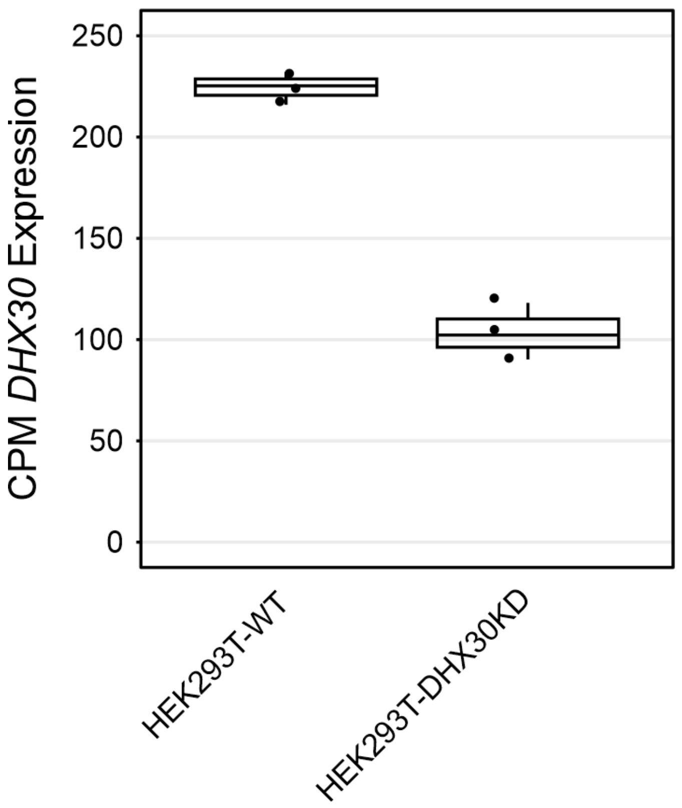


**Supplementary Fig. S16. DHX30 expression in HEK293T cells.** Box plot showing DHX30 expression levels, expressed as counts per million (CPM), in HEK293T wild-type (WT) and DHX30 knockdown (DHX30KD) cells. Boxes represent the interquartile range, the center line indicates the median, and points represent individual replicates. Data presented are from three independent biological experiments.

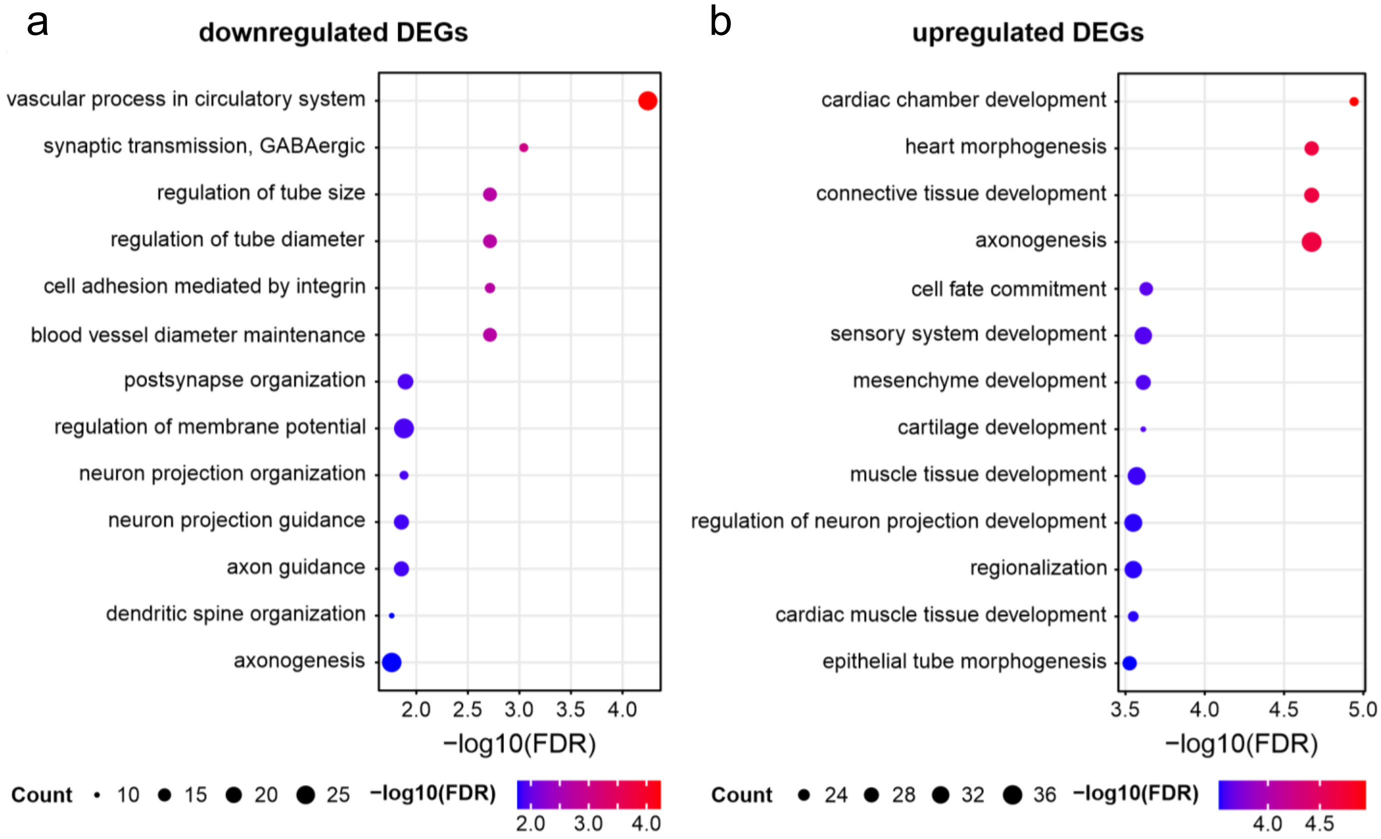


**Supplementary Fig. S17. Gene Ontology enrichment analysis of downregulated and upregulated genes following DHX30 depletion. a** Gene Ontology (GO) enrichment analysis of significantly downregulated differentially expressed genes (DEGs) in HEK293T-DHX30KD cells. **b** GO enrichment analysis of significantly upregulated DEGs in HEK293T-DHX30KD cells. In both panels, dot size indicates the number of genes associated with each GO term, while colour indicates statistical significance (−log10 FDR).

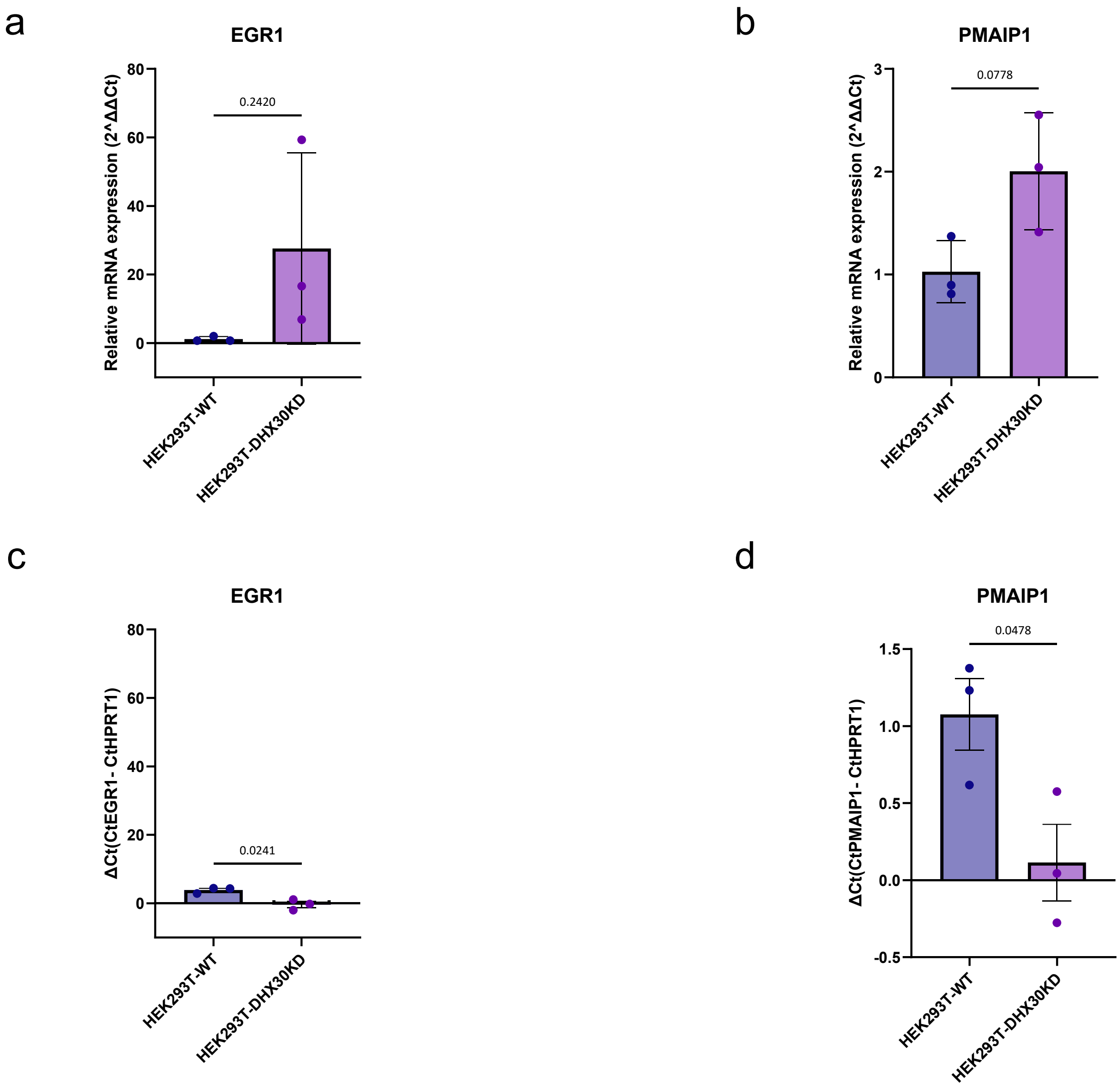


**Supplementary Fig. S18. RT-qPCR validation of EGR1 and PMAIP1 expression changes following DHX30 depletion. a-b** Relative mRNA expression of **EGR1 a** and **PMAIP1 b** in HEK293T-WT and HEK293T-DHX30KD cells, determined by qRT-PCR and expressed as 2^−ΔΔCt. **c-d** Corresponding ΔCt values for EGR1 **(c)** and PMAIP1 **(d)** used for statistical analysis. Data are presented as mean ± SEM from three independent biological experiments. Statistical significance was assessed on ΔCt values using an unpaired two-tailed Welch's t-test (EGR1, $P = 0.0241$; PMAIP1, $P = 0.0478$; $^{*}P < 0.05$).
