## Supplementary table 2 for "DHX30 is a replication stress-responsive R-loop helicase that safe-guards genome stability"

Supplementary Table 2. DNA and RNA radio labelled oligonucleotide used for R-loop and D-loop substrates.

| Name | Sequence 5’-3’ | Reference |
|---|---|---|
| R5_DNA | AACGACGGCCAGTGAATTCGAGCTCGGT ACCCAGTCTTAAGCCTTGACTAGTCAGCT TGACTAAGCGATTGACT | (Thakur et al., 2013) |
| R4_DNA | AGTCTTAAGCCTTGACTAGTCAGCTTGAC TAAGCGATTGACTAACGACGGCCAGTGAA TTCGAGCTCGGTACCC | (Thakur et al., 2013) |
| R3_DNA | AACGACGGCCAGTGAATTCGAGCTCGGT ACCC | (Thakur et al., 2013) |
| T1_DNA | TGTGGAATGCTACAGGCGTTGTAGTTT GTACTGGTGACGAAACTCAGTGTAACG GTACATGGGTTCCTATTGGGCTTGCTATCCCT GAAAATGAGGGTGG | (Thakur et al., 2013) |
| B11_DNA | CCACCCTCATTTTCAGGGATAGCAAGCCCAAT AGGGGTACCGAGCTCGAATTCACTGGCCGTC GTTCCAGTACAAACTACAACGCCTGTAGCATT CCACA | (Thakur et al., 2013) |
| R5_RNA | AACGACGGCCAGUGAAUUCGAGCUCGGUAC CCAGUCUUAAGCCUU | (Thakur et al., 2013) |
| R4_RNA | UAAGCGAUUGACUAACGACGGCCAGUGAAU UCGAGCUCGGUACCC | (Thakur et al., 2013) |
| R3_RNA | AACGACGGCCAGUGAAUUCGAGCUCGGUAC CC | (Thakur et al., 2013) |
