## Supplementary table 3 for "DHX30 is a replication stress-responsive R-loop helicase that safe-guards genome stability"

Supplementary Table 3. DNA and RNA oligonucleotides and schemes of RNA/DNA substrates used for electrophoretic mobility shift assays. These substrates and sequences were previously described Boleslavska, Oravetzova et al. 2022.

| NAME | LENGTH | SEQ. (5´→3´) |
|---|---|---|
| R1-A | 25 | (FAM) - AGCUAUGACCAUGAUUACGAAUUGC |
| R2-R | 35 | (FAM)-AGCUAUGACCAUGAUUACGAAUUGCAAAAAUUUUU |
| R3-N | 35 | AAAAAUUUUU AGCUAUGACCAUGAUUACGAAUUGC- (FAM) |
| D1-B | 25 | (FAM) - AGCTATGACCATGATTACGAATTGC |
| D2-C | 25 | GCAATTCGTAATCATGGTCATAGCT |
| D3-I | 65 | GGGTGAACCTGCAGGTGGGCTTTTTTTTTTTTTTTTTTTTTTTTTGGTAGAATTCGGCAGCGTC |
| D4-J | 65 | GACGCTGCCGAATTCTACCAGCAATTCGTAATCATGGTCATAGCTGCCCACCTGCAGGTTCACCC |

| Name | Scheme |
|---|---|
| Blunt ended RNA:DNA hybrid | 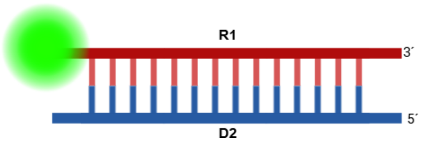 |
| R-loop | 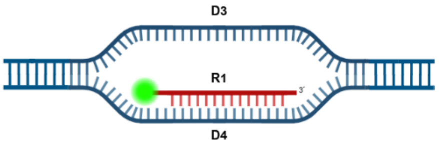 |
| R-loop 10-nt 3’-overhang | 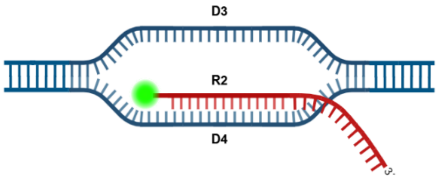 |
| R-loop 10-nt 5’-overhang | 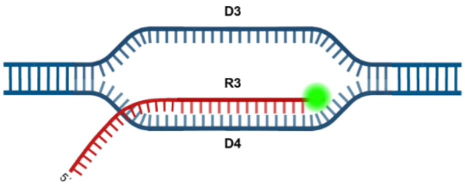 |
| D-loop | 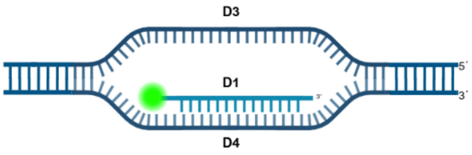 |
